# Fc mediated pan-sarbecovirus protection after alphavirus vector vaccination

**DOI:** 10.1101/2022.11.28.518175

**Authors:** Lily E. Adams, Sarah R. Leist, Kenneth H. Dinnon, Ande West, Kendra L. Gully, Elizabeth J. Anderson, Jennifer F. Loome, Emily A. Madden, John M. Powers, Alexandra Schäfer, Sanjay Sarkar, Izabella N. Castillo, Jenny S. Maron, Ryan P. McNamara, Harry L. Bertera, Mark R. Zweigert, Jaclyn S. Higgins, Brea K. Hampton, Lakshmanane Premkumar, Galit Alter, Stephanie A. Montgomery, Victoria K. Baxter, Mark T. Heise, Ralph S. Baric

**Affiliations:** Department of Microbiology & Immunology, University of North Carolina at Chapel Hill, Chapel Hill, NC, USA; Department of Epidemiology, University of North Carolina at Chapel Hill, Chapel Hill, NC, USA; Division of Comparative Medicine, University of North Carolina at Chapel Hill, Chapel Hill, NC, USA; Department of Genetics, University of North Carolina at Chapel Hill, Chapel Hill, NC, USA; Department of Pathology and Laboratory Medicine, University of North Carolina at Chapel Hill, Chapel Hill, NC, USA; Rapidly Emerging Antiviral Drug Discovery Initiative, University of North Carolina at Chapel Hill, Chapel Hill, NC, USA; Ragon Institute of MGH, MIT, and Harvard University, Cambridge, MA, USA; Dallas Tissue Research, Dallas, TX, USA

## Abstract

Two group 2B β-coronaviruses (sarbecoviruses) have caused regional and global epidemics in modern history. The mechanisms of cross protection driven by the sarbecovirus spike, a dominant immunogen, are less clear yet critically important for pan-sarbecovirus vaccine development. We evaluated the mechanisms of cross-sarbecovirus protective immunity using a panel of alphavirus-vectored vaccines covering bat to human strains. While vaccination did not prevent virus replication, it protected against lethal heterologous disease outcomes in both SARS-CoV-2 and clade 2 bat sarbecovirus HKU3-SRBD challenge models. The spike vaccines tested primarily elicited a highly S1-specific homologous neutralizing antibody response with no detectable cross-virus neutralization. We found non-neutralizing antibody functions that mediated cross protection in wild-type mice were mechanistically linked to FcgR4 and spike S2-binding antibodies. Protection was lost in FcR knockout mice, further supporting a model for non-neutralizing, protective antibodies. These data highlight the importance of FcR-mediated cross-protective immune responses in universal pan-sarbecovirus vaccine designs.

## INTRODUCTION

Three β-coronaviruses (βCoVs) have caused epidemic and pandemic disease in human populations in the 21^st^ century. The 2002-2004 severe acute respiratory coronavirus (SARS-CoV) and the 2019 SARS-CoV-2 (the causative agent of the COVID-19 pandemic) viruses represent prototype clade 1a and clade 1b group 2B βCoVs, which belong to the subgenus sarbecoviruses. SARS-CoV and SARS-CoV-2 likely emerged from bat reservoirs either through intermediate host transmission events or through direct spread into human populations (1, 2). The subgenus sarbecovirus includes other highly heterogeneous epidemic, pandemic, and zoonotic strains poised for emergence and cross-species transmission to novel mammalian hosts. Other sarbecoviruses identified in bat or mammalian reservoirs, including SHC014 and WIV1, can use human ACE2 receptors for entry and replicate efficiently in primary human cells (3–11). Due to global efforts, safe and effective vaccines against SARS-CoV-2 are approved for use (12–14) and next-generation vaccines including those vectored by alphaviruses are under development (15–17). The accelerated approval of SARS-CoV-2 vaccines is the result of decades of basic and applied research (12). However, current SARS-CoV-2 spike-based vaccines provide limited protection against heterologous bat sarbecoviruses, as well as recently emerged SARS-CoV-2 variants of concern (VOC) (18). The extent and complex immunologic mechanisms regulating cross protection against closely related and distant strains remains poorly understood but is critical for pan-coronavirus vaccine design, human health, and public health preparedness.

The sarbecovirus spike glycoprotein is a class I viral fusion protein roughly 1,300 amino acids in length that trimerizes upon folding. Spike is divided into an amino-terminal S1 subunit, containing the receptor-binding domain, and a carboxy-terminal S2 subunit, driving membrane fusion. The S1 subunit is further subdivided into a highly variable N-terminal domain (NTD) and a receptor binding domain (RBD), which engages the ACE2 receptor. Subtle molecular communication networks across domains are thought to influence epitope presentation, as evidenced by druggable targets in the NTD that interrupt distal RBD interactions with ACE2 as well as the identification of spike mutations outside of the RBD that stabilize receptor interaction (19, 20). As such, structural features of the spike are likely to impact vaccine cross-protection. Recent work has characterized the immune response against distinct Sarbecovirus spike proteins following homologous or heterologous vaccination (21–24). Cross-reactive T cells and antibodies recognize broadly conserved epitopes across SARS-CoV-2, other sarbecoviruses, and endemic (common-cold) β-coronaviruses. However, the role of these epitopes in protective immunity remains a subject of rigorous investigation (22, 25, 26). After SARS-CoV-2 natural infection or vaccination, the spike RBD, NTD, and S2 domains stimulate neutralizing and non-neutralizing antibody responses. Among sarbecoviruses, potent, broadly protective, neutralizing antibodies primarily target specific epitopes in the RBD and S2 portions of the spike glycoprotein (23, 27–32), typically targeting epitope bins RBD-6 and RBD-7 (33, 34). However, neutralizing antibody activity and function have not been correlated, as it is sometimes difficult to predict whether potent cross-reactive neutralizing antibodies that target spike will sufficiently protect *in vivo* (34). Although less widely studied, several studies have shown that antibody FcR-mediated effector functions are critical in protective immunity (35–39). Fc receptors have high affinity for IgG subtypes and are cell surface receptors on monocytes, macrophages, and neutrophils; FcR recognition of the antibody Fc region stimulates effector cell function like NK cell-mediated lysis, neutrophil degranulation, and ADCP (40). Thus, FcR effector function may be a key correlate for next generation vaccine development and improvement.

Here, we utilized a Venezuelan equine encephalitis virus 3526 replicon particle (VRP3526) (41) as a platform to evaluate mechanisms of cross protection after pandemic and pre-emergent coronavirus spike glycoprotein vaccination in mice, followed by lethal SARS-CoV-2 challenge (42). VRP vectors induce robust cellular and humoral immune responses after vaccination and VRP spike vaccines protect against SARS-CoV and SARS-CoV-2 *in vivo* (43, 44). In the present study, contemporary human coronavirus spikes elicited no protection against weight loss, mortality, or virus replication after SARS-CoV-2 challenge. While VRP delivered homologous SARS-CoV-2 spike vaccines protected against weight loss, lethal disease, and virus replication after homologous challenge, heterologous VRP sarbecovirus spike vaccines conferred cross protection against weight loss and death, but provided limited reductions in SARS-CoV-2 MA10 replication in young and aged animals. While homologous protection correlated with potent neutralizing antibody responses that principally targeted the S1 subdomains, no cross-neutralizing antibodies were detected against the heterologous sarbecovirus strains. Rather, systems serology (45), *in vitro* studies, and passive antibody transfer experiments in wild-type and Fc-receptor deficient mice implicated an FcR-driven mechanism targeting S2, such as antibody-dependent cellular phagocytosis (ADCP). These results build support for universal sarbecovirus vaccine designs that include FcR-mediated cross protection, coupled with potent cross-neutralizing antibody responses.

## RESULTS

### Venezuelan equine encephalitis virus replicon particle assembly produced high titer vaccines to potently express coronavirus spike proteins *in vivo*

Venezuelan equine encephalitis virus 3526 replicon particles (VRPs) are BSL2, non-select, replication-deficient vectors derived from a live attenuated strain (41). They are assembled using a tripartite RNA-based assembly scheme (**Fig. 1A**), thus generating a self-amplifying, single cell hit RNA vaccine platform. We generated replicon particles (VRPs) expressing spike proteins from several α- and β-CoVs that included three common-cold CoVs, OC43, NL63, and HKU1 (**Fig. 1B, green**), as well as pandemic (SARS-CoV, SARS-CoV-2) and pre-emergent sarbecoviruses circulating in animal reservoirs (RaTG13, HKU3, WIV1, SHC014) (**Fig. 1B, red**). The sarbecovirus spike proteins were separated into three groups based on amino acid similarity: clade 1a (SARS-CoV, SHC014, WIV1), 2 (HKU3), and 1b (SARS-CoV-2, RaTG13) (**Fig. 1A**) (46). The clade 1b virus RaTG13 spike protein is 97.4% identical to the SARS-CoV-2 spike protein, the clade 2 virus HKU3 spike protein shares 75.8% identity to the SARS-CoV-2 spike protein, and the clade 1a virus spike proteins share 75.6-78.6% identity to the SARS-CoV-2 spike protein. Clade 2 HKU3 spike protein shares 78.1-78.8% identity to the clade 1a spike proteins (**Table 1**). All VRP preparations achieved particle titers exceeding 2×10^6^ IU/mL, sufficient for vaccination in a mouse model (**Fig. 1C)** (41). Immunofluorescent staining for the highly conserved spike S2 domain verified spike expression in mammalian cells infected by VRPs (**Fig. 1D**).

**Figure 1.**
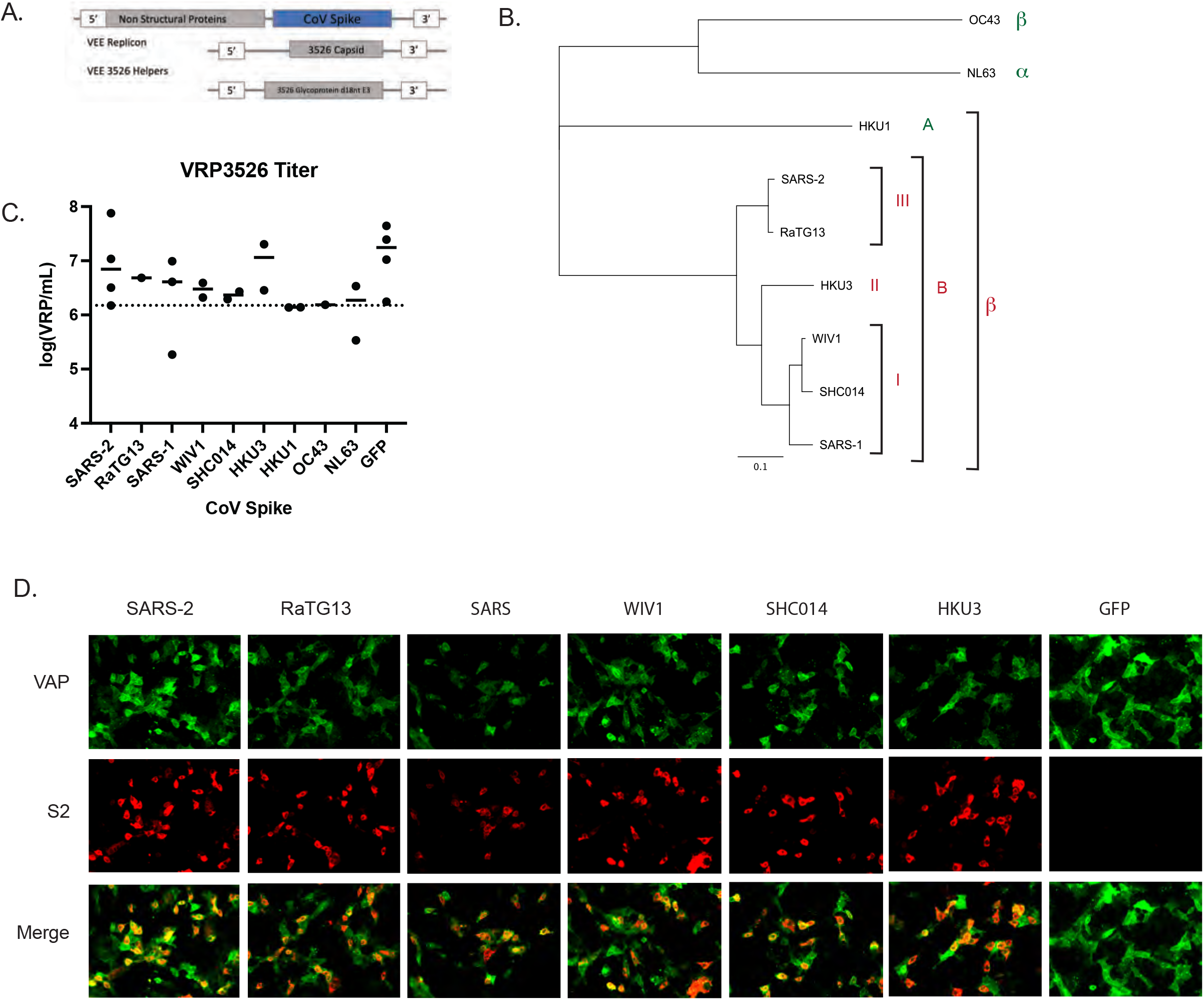
Venezuelan Equine Encephalitis Virus Replicon Particle VRP3526 for high-titer vaccinations. **A.** Fragmented RNA-based assembly scheme of VRP3526 particles. **B.** Phylogenetic relationships of CoV spike proteins that were used in this study, including common cold CoVs (green) and prepandemic/epidemic CoVs (red). Of the β-coronaviruses, we generated spike proteins for both group 2A (HKU1) and 2B viruses. Of the group 2B viruses, we generated spike proteins for clade 1a (SARS-CoV, SHC014, WIV1), 2 (HKU3), and 1b (SARS-CoV-2, RaTG13) viruses. Tree generated from an amino acid multiple sequence alignment using Maximum Likelihood in Geneious Prime. **C.** VRP3526 titers obtained in this study. Dashed line denotes minimum titer required for vaccination at 2×10^4^ VRP in a 10 μl footpad inoculation. **D.** Immunofluorescent staining at 40x magnification for VEE non-structural proteins (top) and SARS-CoV-2 spike S2 domain (middle) in Vero E6 cells infected with VRPs expressing the spike proteins used in this study.

**Table 1.** Amino acid percent similarity | patristic phylogenetic distance of sarbecovirus spike proteins

|  | SARS-CoV-2 | RaTG13 | SARS-CoV | SHC014 | WIV1 |
| --- | --- | --- | --- | --- | --- |
| RaTG13 | 97.4 \| 0.023 |  |  |  |  |
| SARS-CoV | 75.6 \| 0.248 | 76.2 \| 0.247 |  |  |  |
| SHC014 | 76.8 \| 0.246 | 77.0 \| 0.245 | 90.0 \| 0.099 |  |  |
| WIV1 | 76.5 \| 0.233 | 77.0 \| 0.232 | 92.3 \| 0.086 | 97.1 \| 0.030 |  |
| HKU3 | 75.8 \| 0.248 | 76.0 \| 0.247 | 78.1 \| 0.225 | 78.8 \| 0.223 | 78.8 \| 0.210 |

### VRP-vectored sarbecovirus spike proteins protect against severe SARS-CoV-2 disease in young mice

To test the VRP 3526 platform and evaluate the capacity for spike protein elicited cross-protection, we utilized a lethal mouse model for SARS-CoV-2 disease. Groups (n = 8-10) of 8-10 week aged female BALB/cAnNHsd (BALB/c) mice were vaccinated with a low dose of 2×10^4^ IU VRP encoding each of the different spike vaccines by footpad injection then boosted on day 21 with the homologous spike VRP. At 21 days post-boost, (now 14-16 week aged) BALB/c mice were challenged with 10^4^ PFU SARS-CoV-2 MA10 (42) intranasally. Virus titer after challenge is a sensitive measure of vaccine performance, and reductions in titer are often correlative to vaccine efficacy (12, 47). However, only the VRP SARS-CoV-2 spike vaccine elicited nearly complete protection from homologous virus replication. In contrast, the clade 1b VRP RaTG13 spike vaccination resulted in slight, but significant reductions in virus titers on day 2 (1 log reduction, 10-fold) and 5 (3 log reduction, 1,000-fold) post infection, as compared to the GFP vaccinated controls. In animals vaccinated with clade 2 VRP HKU3 spike, SARS-CoV-2 MA10 titers were significantly reduced by about 1.5 and 3 logs (~30- and 1,000-fold) on days 2 and 5 post infection, compared to GFP control vaccinated animals. We also saw slight, but significant, reductions in virus titers in mice vaccinated with highly heterogeneous clade 1a strains (SARS-CoV, WIV1 and SHC014) on day 2. However, titers were reduced by about 2 logs on day 5 post infection (**Fig. 2A**).

**Figure 2.**
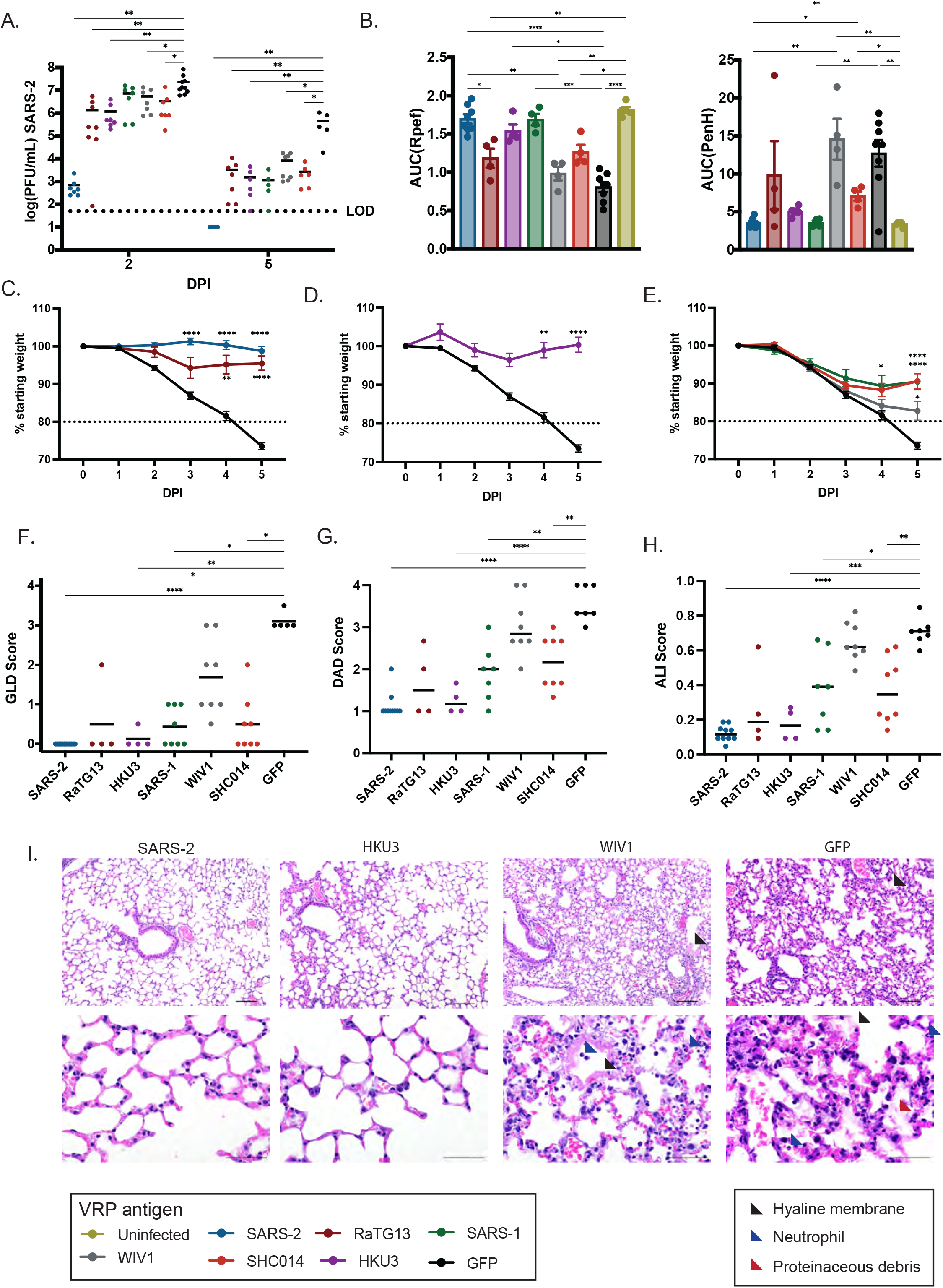
VRP sarbecovirus spike vaccines elicit a cross-protective immune response against SARS-CoV-2. **A** SARS-CoV-2 lung titer calculated via plaque assay on days 2 and 5 post infection. Samples that fell below the limit of detection (dotted line) were set to 25 PFU/mL. **B.** Area under the curve (AUC) of lung function metrics of airflow resistance (PenH, right) and bronchoconstriction (Rpef, left). Lung function was measured by BUXCO whole body plethysmography systems one each experimental day, AUC calculated for time course of each mouse. **C-E.** Body weights calculated after infection of 10^4^ PFU SARS-CoV-2 intranasally through the duration of the experiment on animals vaccinated with clade 1b (**C**), 2 (**D**), and 1a (**E**) sarbecovirus spike proteins. Reported as percent of starting weight. Horizontal line indicates 20% body weight lost and animal care humane endpoint. **F.** Semi-quantitative gross lung discoloration (GLD) scoring, **G.** Diffuse alveolar damage (DAD) scoring, and **H.** acute lung injury (ALI) scoring upon tissue harvest at day 5 post infection. **I.** Hematoxylin and eosin stained sections of lungs from vaccinated mice harvested day 5 post infection. Black arrow – hyaline membrane, blue arrow – neutrophil infiltrate, red arrow – proteinaceous debris. Top – 100x magnification, 100 μm scale. Bottom – 400x magnification, 50 μm scale. * p < 0.05, ** p <0.01, *** p < 0.001, **** p < 0.0001 after statistical testing described in methods.

Consistent with VRP vaccine effects on viral replication, we observed a range of protection outcomes following virus challenge. Bronchoconstriction and airway resistance in the lungs of challenged mice are a representative disease metric measured by whole-body plethysmography that mirrors disease in humans (48). Using this method, we tracked respiratory function through the duration of the experiment and analyzed the overall difference in lung function between vaccine groups via area under the curve calculations. Overall, we found that clade 1b and clade 2 vaccines effectively protected against bronchoconstriction (Rpef) and airway resistance (PenH) after SARS-CoV-2 MA10 challenge **(Fig. 2B**), accordant with reduced clinical disease. In contrast, clade 1a vaccinated animals had significantly increased respiratory dysfunction and clinical disease when compared to uninfected controls, indicating reduced protection. As additional measures of disease severity, we monitored weight loss and assessed lung pathology by scoring gross discoloration (GLD), diffuse alveolar damage (DAD), and acute lung injury (ALI) following SARS-CoV-2 challenge. Homologous SARS-CoV-2 challenge in VRP SARS-CoV-2 spike vaccinated animals resulted in minimal weight loss (**Fig. 2C**) and GLD (**Fig. 2F**), and as such animals were fully protected against significant SARS-CoV-2 disease. In contrast, the other zoonotic and pandemic CoV spike proteins partially protected from clinical disease compared to controls. For example, the clade 1b CoV spike proteins protected against severe SARS-CoV-2 disease, resulting in little (~10%-RaTG13) to no measurable weight loss (SARS-CoV-2) and minimal GLD at the time of tissue harvest (**Fig. 2C, F**). Under identical conditions, clade 1a spike (SARS-CoV, WIV1 and SHC014) vaccines elicited low level intermediate protection after SARS-CoV-2 MA10 challenge, resulting in more weight loss, ranging between 10-15% body weight lost, and notable increases in GLD (**Fig. 2E, F**). Despite being as distant as the clade 1a CoV spikes were from the SARS-CoV-2 spike (Table 1), the clade 2 HKU3 spike vaccine elicited near full protection against disease with ~5% weight loss and minimal lung discoloration at tissue harvest (**Fig. 2D, F**). In addition to antigenic distance, the HKU3 spike protein also contains deletions in both the NTD and RBD when compared to the SARS-CoV-2 spike protein (**Fig. S1**). Thus, the protection elicited by VRP HKU3 spike after SARS-CoV-2 heterologous challenge is particularly noteworthy. Within vaccine groups where mice were partially protected from disease, such as WIV1 spike vaccine, disease outcomes ranged from mild disease (mild weight loss and lung discoloration) to severe disease (significant weight loss and severe lung discoloration), suggesting that this protective mechanism is subject to strain-specific variation, resulting in highly variable disease outcomes. Overall, we calculated a robust negative correlation coefficient (−0.76) between disease metrics of GLD and area under the curve of percent body weight as well as a strong positive correlation (0.72) between virus titers day 2 and day 5 post infection. The slight, but incomplete, reduction in virus titer suggests that protection was likely not mediated by the presence of potent cross-neutralizing antibodies with the ability to prevent infection. However, we calculated a negative correlation coefficient (−0.68) between viral titer two days post infection and disease severity as measure by area under the curve of percent body weight through the duration of the study, indicating virus titer may still be predictive of disease severity in our model.

Consistent with prior established work (42), histological examination (**Fig. 2G, H**) of lung sections stained for hematoxylin and eosin at 5 days post infection identified regions of severe disease in mock vaccinated, infected animals, including infiltration of neutrophils in the interstitial and alveolar spaces, alveolar septal thickening, cell sloughing and proteinaceous debris in the airspaces, and hyaline membrane formation (**Fig. 2I**). Using previously described scoring metrics for diffuse alveolar damage (DAD) and acute lung injury (ALI), histologic sections revealed large numbers of infiltrating immune cells, increased membrane thickness, and proteinaceous debris in the airspaces (42) (**Fig. 2I, red, blue, black**). Consistent with other metrics reported in this study (e.g. weight loss), mice vaccinated with the homologous spike demonstrated protection from SARS-2-induced lung pathology, with baseline-equivalent DAD and ALI scores (41) (**Fig. 2G, H, I**). Heterologous vaccines that were associated with greater GLD scores and thus only partial protection (e.g. WIV1) demonstrated more severe tissue damage and higher DAD and ALI scores. Vaccines that were more protective (e.g. HKU3) resulted in lower DAD and ALI scores comparable to the SARS-2 vaccinated group.

We also used our SARS-CoV-2 mouse lethal challenge model to evaluate whether vaccination with VRPs expressing spikes of contemporary common cold human β-CoV, which share conserved S2 epitopes with epidemic and pre-emergent β-CoV (49, 50) would protect against SARS-CoV-2 disease. We found that single exposures (single component, two doses) of contemporary human coronavirus spike proteins did not protect against severe SARS-CoV-2 disease and mortality in young mice (**Fig. S2 A, C**), nor did these vaccines reduce viral replication efficiency (**Fig. S2 B**). In contrast to the Group 2B coronavirus vaccinated mice, a large percentage (>50% in most cases) of common cold spike vaccinated mice died when compared to those vaccinated with VRP SARS-CoV-2 spike (**Fig. S2 C**).

Evaluating the potential for aberrant immunity after vaccination is especially important as killed/inactivated CoV vaccines have been reported to induce a pro-inflammatory and Th2 skewed immune response, commonly associated with immune pathology (47, 51). Using a BioPlex Cytokine Immunoassay, we measured the cytokine responses after challenge in vaccinated mice on days 2 and 5 post infection. In groups that received heterologous VRP spike vaccines, we did not detect elevated Th2 cell cytokine signatures (IL-4, IL-5, IL-13), rather VRP vaccines elicited a strong Th1 signature (IL-12, TNF-α, IFN-γ) which is commonly associated with a protective immune response (41). Additionally, in contrast to the groups vaccinated with the homologous SARS-CoV-2 spike, groups vaccinated with the heterologous sarbecovirus VRP spikes demonstrated elevated pro-inflammatory cytokine responses in the lung two days post-SARS-CoV-2 MA10 challenge (IL-1β, IFN-γ, TNF-α, IL-6) (**Fig. S3**). Altogether, this suggests the VRP sarbecovirus spike vaccines elicited a protective immune response against SARS-CoV-2 infection.

### VRP-vectored sarbecovirus spike proteins protect against severe SARS-CoV-2 disease in old mice

To test the efficacy of the VRP platform and evaluate the potential for cross-protection in an aging population, we utilized the SARS-CoV-2 MA10 lethal challenge model in aged (1-year old) mice. Mice were immunized with 2×10^4^ IU VRP by footpad inoculation following a prime/boost schedule on days 0 and 21. Three weeks post boost, mice were challenged with 10^3^ PFU (~1 LD_50_, typically 50% mortality) SARS-CoV-2 MA10 intranasally (**Fig. 3A–F**). We observed near complete protection in our homologous vaccinated, aged mouse model, as animals challenged with SARS-CoV-2 MA10 experienced ~5% weight loss before recovery (**Fig. 3A**). Importantly, VRP SARS-CoV-2 spike vaccinated animals were also significantly protected from GLD upon tissue harvest on days 2 and 5 post infection (**Fig. 3D**). Moreover, virus titers were reduced by 5 and 3 logs on day 2 and 5 post infection, a significant reduction when compared to controls (**Fig. 3E**). In contrast, clade 1b VRP RaTG13 spike vaccinated animals showed some protection from clinical disease as measured by limited weight loss (~8%) and GLD on days 2 and 5 post infection, as compared to mock vaccinated controls (**Fig. 3A, D**). Clade 2 VRP HKU3 spike vaccinated animals also showed limited weight loss (~10%), GLD, and slight reduction in virus titer on days 2 and 5 post infection, which were significantly different from controls. Clade 1a VRP spike vaccines elicited significant but variable levels of protection in the aged mouse model after SARS-CoV-2 MA10 challenge (**Fig. 3B, D**). Based on protection from weight loss and lung discoloration scores, clade 1a VRP vaccines were protective when compared to mock vaccinated controls on days 2 and 5 post infection (**Fig. 3C, D**). However, clade 1a vaccines did not protect against virus replication on day 2, as modest but not significant ~1 log reductions were noted in VRP WIV1 and VRP SHC014 vaccinated animals on day 5. Reductions in respiratory function generally correlated with overall disease severity, as measured by weight loss and GLD (**Fig. 3F**). Compared to young mice, SARS-CoV-2 replicated to higher titers in the lungs of the old mice with more breakthrough replication in the homologous vaccinated mice (**Fig. 3E**).

**Figure 3.**
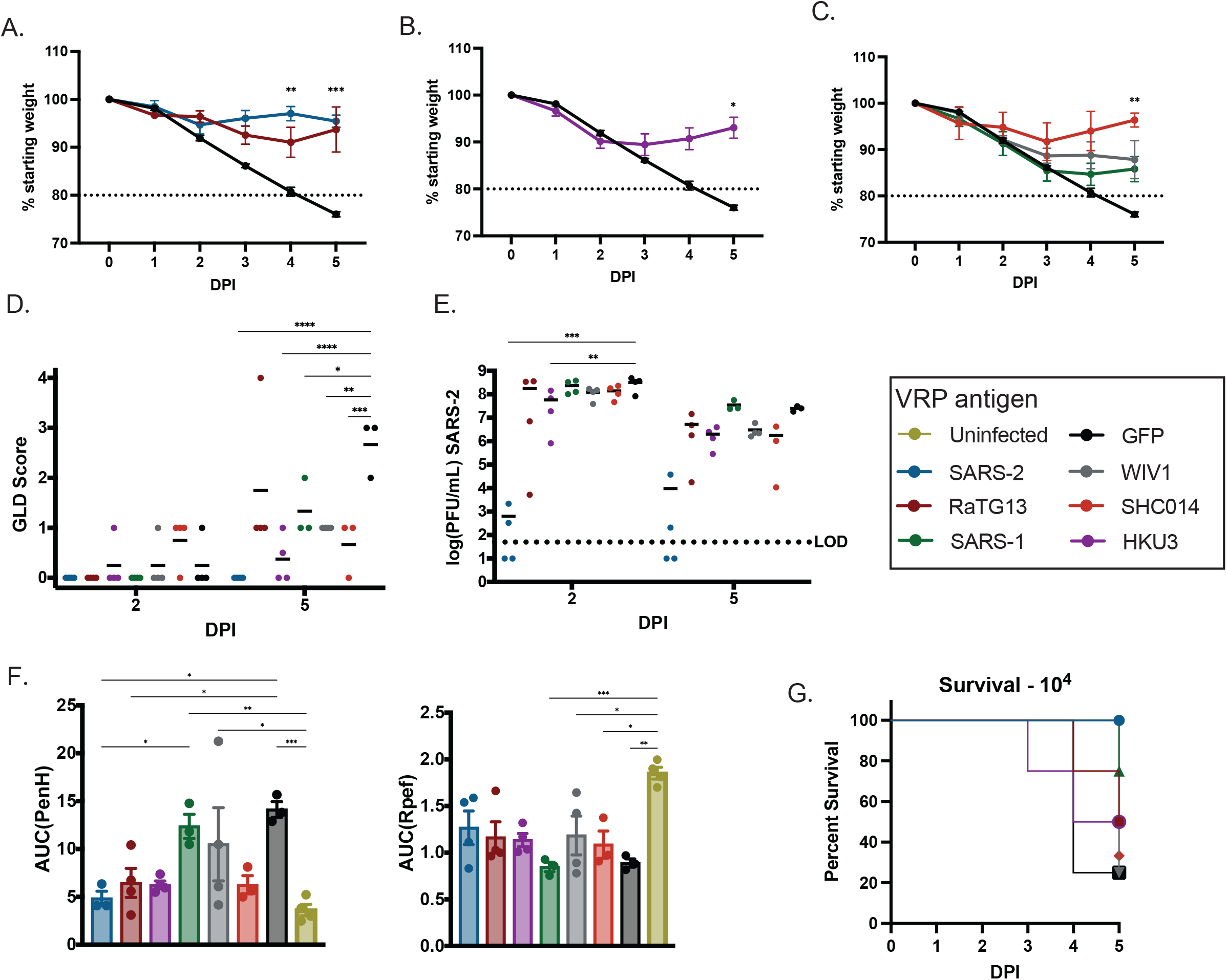
VRP Spike protects from lethal infection in vulnerable aged mice. Old mice (12 months) were challenged with 10^3^ PFU SARS-CoV-2 MA10 intranasally unless otherwise noted. **A-C.** Body weights calculated through the duration of the experiment on animals vaccinated with clade 1b (**A**), 2 (**B**), and 1a (**C**) sarbecovirus spike proteins. Reported as percent of starting weight. Horizontal line indicates 20% body weight lost and animal care humane endpoint. **D.** GLD scoring upon tissue harvest. **E.** SARS-CoV-2 lung titer calculated via plaque assay. Samples that fell below the limit of detection (dotted line) were set to 25 PFU/mL. **F**. Area under the curve (AUC) of lung function metrics of airflow resistance (PenH, right) and bronchoconstriction (Rpef, left). Lung function was measured by BUXCO whole body plethysmography systems one each experimental day, AUC calculated for time course of each mouse. **G.** Survival of vaccinated animals when challenged with 10^4^ PFU SARS-CoV-2 MA10 intranasally. * p < 0.05, ** p <0.01, *** p < 0.001, **** p < 0.0001 after statistical testing described in methods.

As aged animals are significantly more vulnerable to higher SARS-CoV-2 MA10 challenge doses (42), we next determined if the VRP sarbecovirus spike vaccine panel could protect against a 10-fold higher, 10^4^ challenge dose, which typically results in 85% mortality. Mice vaccinated with VRP SARS-CoV-2 spike were fully protected from severe disease and mortality through day 5 post infection. In contrast, mice vaccinated with other clade 1a, 1b, or clade 2 heterologous VRP spike vaccines experienced equivalent weight loss as mock vaccinated controls (**Fig. S4**) and mortality rates of 25-75% (**Fig. 3G**), indicating the cross-protection elicited in our model has dramatic potential to vary in an aging population, especially as a function of infectious dose.

### VRP SARS-CoV-2 spike protects against disease in heterologous sarbecovirus infection

The clade 2 HKU3 strain cannot infect primate cells and does not use an ortholog human, civet, select bat, or mouse ACE2 receptor for docking and entry (11). However, more recent studies suggest that some clade 2 strains can utilize bat ACE2 molecules for entry if isolated from their natural bat host species (52), which suggests clade 2 sarbecoviruses may become an emergent threat in the future. Although the HKU3 spike is phylogenetically distant from clade 1 strains (**Table 1**), vaccination elicited a good protective profile against SARS-CoV-2 MA10 (**Fig. 2**). As HKU3 could emerge by mutation or RNA recombination, we next evaluated whether the VRP SARS-CoV-2 spike vaccine would protect against clade 2 heterologous challenge. 8-10 week old mice were vaccinated and boosted with VRP HKU3, VRP SARS-CoV-2, and VRP GFP vaccines as previously described. Vaccinated mice were then infected intranasally with the mouse-adapted clade 2 bat sarbecovirus designated HKU3-SRBD MA, a virus that can infect mammalian cells and cause disease in mice (11, 53). The homologous HKU3 vaccinated animals were fully protected from weight loss, GLD, and showed a significant 4-log reduction in titer after HKU3-SRBD challenge (**Fig. 4A–C**). In contrast, the heterologous VRP SARS-CoV-2 vaccine attenuated HKU3-SRBD disease severity as evidenced by ~15% body weight loss (**Fig. 4A**), a recovery of body weight after 3 days, and modest reductions in GLD scores when compared to mock vaccinated animals (**Fig. 4B**). Modest but significant ~5- and 10-fold reductions in virus titer were also noted on days 2 and 5 post infection, respectively, in VRP SARS-CoV-2 vaccinated mice when compared to VRP GFP vaccinated control (**Fig. 4C**). As such, we observed a cross-protective phenotype mediated by VRP spike vaccines in two unique challenge models, prompting further mechanistic investigation.

**Figure 4.**
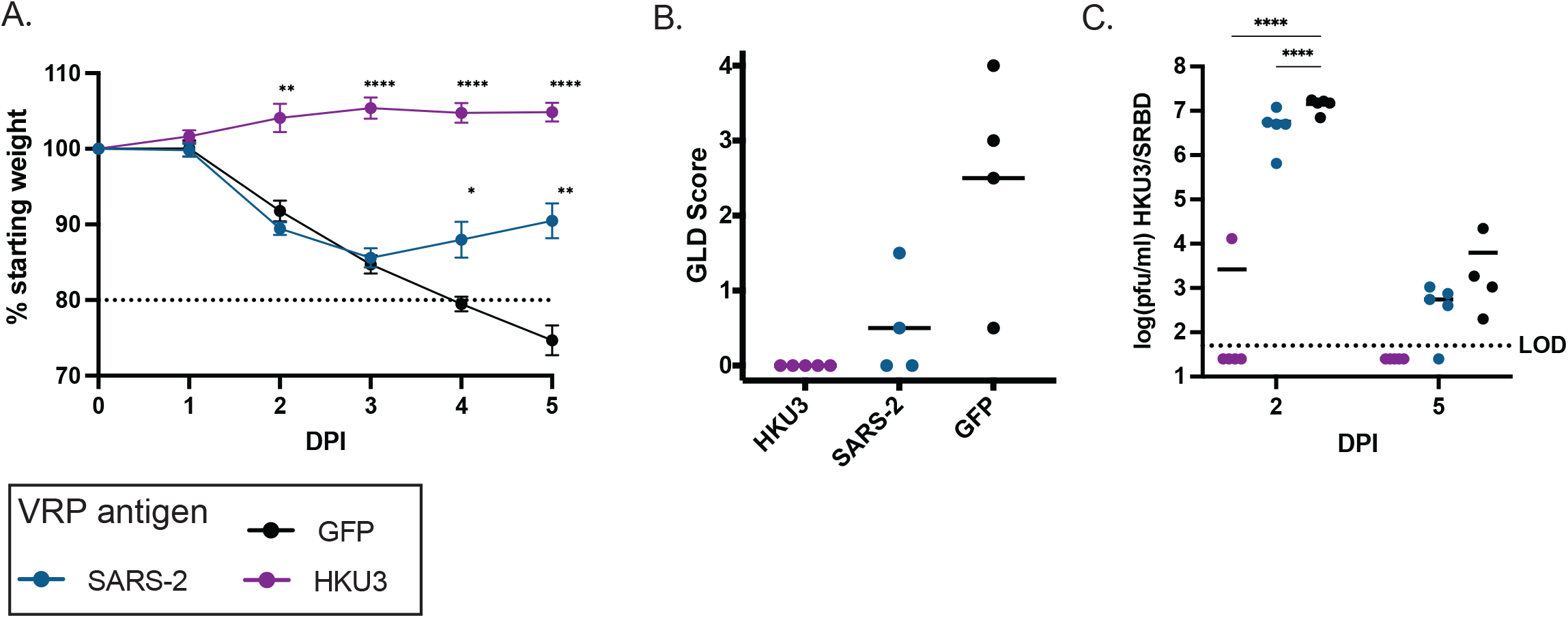
VRP SARS-CoV-2 spike vaccination protects against heterologous challenge. Young mice (16-18 weeks) were challenged with 10^5^ PFU HKU3/sRBD MA (**A**-**C**) or old mice (12 months) were challenged with 10^5^ PFU SARS-CoV-2 MA10 containing the omicron variant spike (**D**-**F**) intranasally. **A.** Body weights calculated through the duration of the experiment on animals vaccinated with SARS-CoV-2 or HKU3 spike proteins, or vectored GFP. **B.** Semi-quantitative macroscopic lung discoloration scoring upon tissue harvest. **C.** HKU3 lung titer calculated via plaque assay. * p < 0.05, ** p <0.01, *** p < 0.001, **** p < 0.0001 after statistical testing described in methods. Body weights reported as percent of starting weight where horizontal line indicates 20% body weight lost and animal care humane endpoint. Titer samples that fell below the limit of detection (dotted line) were set to 25 PFU/mL.

### VRP spike vaccinations induce non-neutralizing, cross-reactive antibodies

Each VRP spike vaccine elicited a potent serologic IgG response against the SARS-CoV-2 spike protein (**Fig. 5A, left**). Total IgG titers ranged from 1 x 10^4^ to 5 x 10^4^. The VRP spike vaccines also elicited a potent IgG response against the SARS-CoV-2 receptor binding domain (RBD**, Fig. 5A, middle**), though at about a two-fold reduction in potency. Additionally, mice vaccinated with VRP RaTG13 did not produce significant IgG against the SARS-CoV-2 RBD, despite inducing notable IgG against the SARS-CoV-2 N-terminal domain, comparable to the homologous VRP SARS-CoV-2 vaccine (NTD, **Fig. 5A, right**). The NTD is fairly well conserved between SARS-CoV-2 and RaTG13 (98.3% amino acid identity), while the RBD is more diverged (89.3%, more divergence in the receptor-binding motif), so these results were not unexpected. After VRP vaccination (**Fig. 5B**), we detected a strong IgG2a skew in antibody titers in VRP spike antigen-vaccinated mice, but not in GFP control vaccinated mice, indicating that vaccination with the sarbecovirus spikes induced a protective Th1 response (54). We also observed a marked vaccine antigen specific grouping in the ratio of IgG2a to IgG1; for example, the VRP SARS-CoV-2 vaccine (blue) elicited a more IgG2a skewed antibody response against the SARS-CoV-2 spike protein than the VRP RaTG13 vaccine (red). Further analyses demonstrated high reactivity towards full-length spike with little-to-no preference for IgG2a recognition of S1 or S2 (**Fig. 5C**). Given the presence of cross-reactive antibody responses elicited by VRP-spike vaccines, we then tested the neutralizing antibody response in a series of live-virus assays. Overall, we detected neutralizing antibodies only in the homologous VRP SARS-CoV-2 vaccine group with an IC_50_ of ~1:700 but did not detect any cross-neutralizing antibody responses between any of the other VRP vaccine groups and luciferase reporter SARS-CoV-2, except for some low-level SARS-CoV-2 neutralizing titers in a subset of VRP RaTG13 vaccinated animals (**Fig. 5D**). Additionally, VRP SHC014 sera neutralized reporter virus SHC014 with an IC_50_ of ~1:800 (**Fig. 5F**), but there was also no detectable neutralization of the SHC014 virus by VRP SARS-CoV-2 sera.

**Figure 5.**
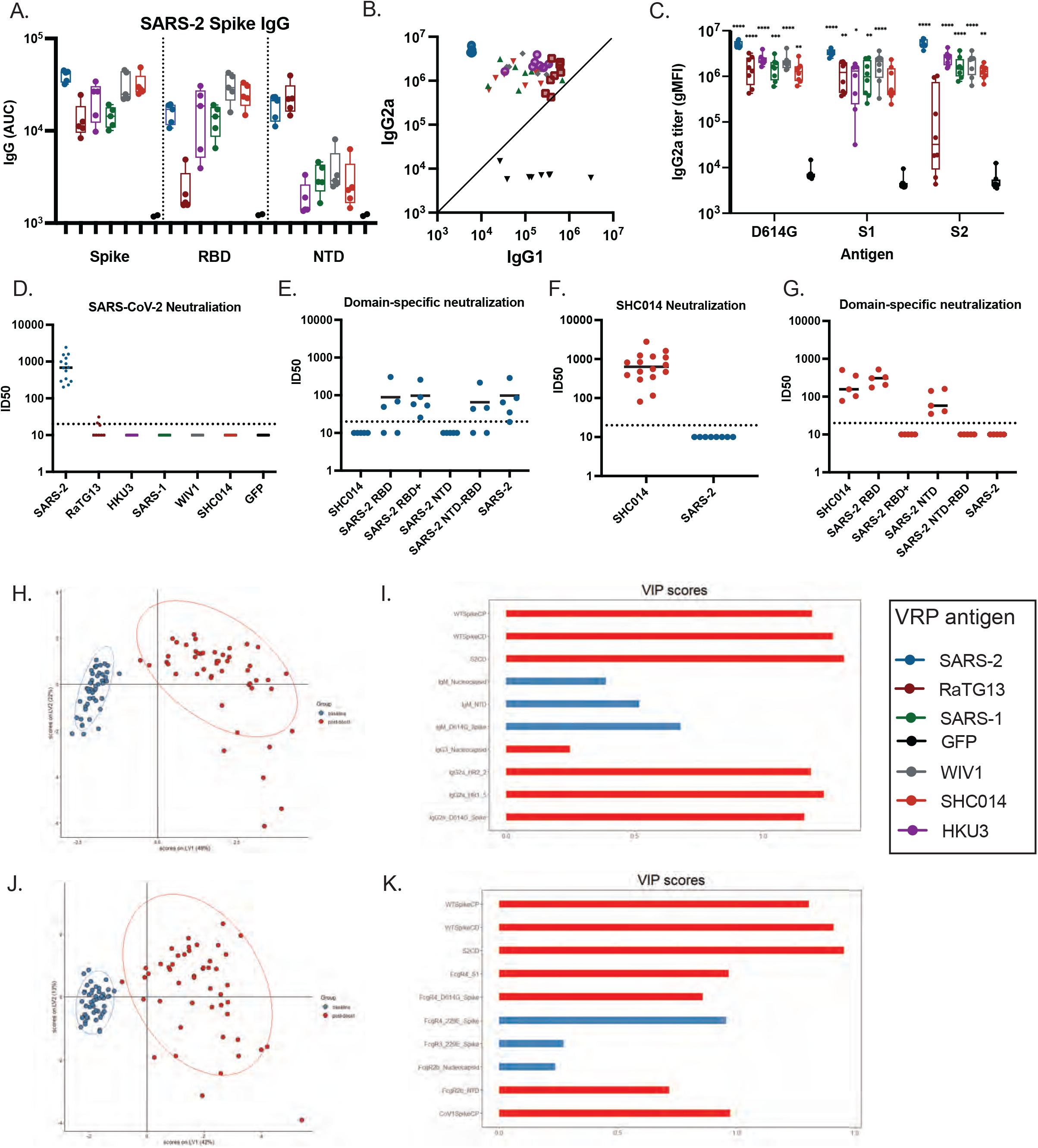
Characterizing the antibody response through systems serology and functional assays. **A.** SARS-CoV-2 spike (left), receptor binding domain (middle), and N-terminal domain (right) binding IgG quantified by ELISA. Titers calculated via area under the curve (AUC). **B.** SARS-CoV-2 spike binding IgG2a titers plotted against IgG1 titers. Titer calculated by gMFI. **C.** SARS-CoV-2 spike (left), S1 (middle), and S2 (right) binding IgG2a titer calculated by gMFI via Luminex bead assay. * p < 0.05, ** p <0.01, *** p < 0.001, **** p < 0.0001 after statistical testing described in methods, comparing each vaccine group to GFP. **D.** SARS-CoV-2 neutralization IC_50_ values calculated as the serum dilution that achieved 50% neutralization in a live-virus neutralization assay. **E.** Domain-specific IC_50_ values measured by live-virus neutralization assay against different SARS-CoV-2 spike domains on a heterologous virus backbone (SHC014). **F.** SHC014 neutralization IC_50_ values of serum from vaccinated animals. **G.** Domain-specific IC_50_ values measured by live-virus neutralization assay against different SARS-CoV-2 spike domains on the SHC014 backbone. IC_50_ for samples that fell below the limit of detection (dotted line) were set to 10. **H.** A scores plot representing the baseline (blue) and post-boost (red) vaccine immunoglobulin and functional profile distribution for all vaccinated animals tested, clustered via PLSDA (partial least squares discriminant analysis). **I.** VIP score of most influential features, representing the total distance from the center of the scores plot, as determined by PLSDA of immunoglobulin and functional profiles. **J.** A scores plot representing the baseline (blue) and post-boost (red) vaccine Fc receptor stimulation and functional profile distribution for all vaccinated animals tested, clustered via PLSDA (partial least squares discriminant analysis). **K.** VIP score of most influential features, representing the total distance from the center of the scores plot, as determined by PLSDA of Fc receptor stimulation and functional profiles.

Using a luciferase reporter system to detect spike NTD, RBD and S1 domain-specific neutralizing antibodies (18, 34), we determined that the majority of VRP SARS-CoV-2 spike neutralizing antibodies targeted the RBD (amino acids 332-528) and the C-terminal segment of S1 (RBD+, amino acids 332-685). Additionally, despite clear evidence of at least 4 neutralizing epitopes in the N-terminal domain (NTD, amino acids 13-305) (33), the VRP SARS-CoV-2 spike vaccines failed to elicit measurable neutralizing antibody titers against the SARS-CoV-2 NTD or S2 domain (**Fig. 5E**). We also mapped the domain specific neutralizing antibody responses elicited by animals vaccinated with the VRP SHC014 spike. In this loss of function assay, VRP SHC014 spike elicited neutralizing antibodies preferentially targeted the NTD and RBD+ regions. The complete loss of neutralization was noted when larger segments of the spike protein were exchanged in the reporter system – either RBD+ or the region spanning from the beginning of the NTD to the end of the RBD (NTD-RBD, amino acids 13-528). This also indicates that SHC014 homologous neutralizing antibodies preferentially target highly specific epitopes in the SHC014 spike S1 region (**Fig. 5G**). Our data suggests that neutralizing antibody responses elicited by VRP spike vaccines are highly type-and domain-specific, supporting the hypothesis that cross-neutralizing antibodies do not drive VRP cross-protection between sarbecoviruses.

Given the inconsistency between neutralizing antibodies and cross protection, we further employed systems serology to characterize the overall humoral architecture in response to our VRP candidates (37, 45) (**Fig. S5**). An initial multivariate analysis (partial least squares discriminant analysis, or PLS-DA) demonstrated a strong clustering of challenged animals away from baseline for both Fab (**Fig. 5H**) and FcR (**Fig. 5J**) binding antibodies. Individual features that separated groupings were identified. Interestingly, Fc-mediated, non-neutralizing functions such as antibody dependent complement deposition (ADCD) and antibody-dependent cellular phagocytosis (ADCP) were among the highest ranked. We validated this through clustering these functional assays with both Fab and FcR binding profiles (**Fig. 5I, K**). Similar to our previous analysis, PLS-DA identified that humoral recognition of both S1 and S2 subregions was driving the phenotype, and not simply RBD-responsive antibodies, which bear the majority of neutralizing activity.

To more closely delineate protective signatures stimulated by the VRP spike vaccines, we performed cross-correlative analyses for the entire data set as well as each VRP spike vaccine (**Fig. S6**). To summarize the correlation matrices and identify trends, we highlighted associations with strong, significant correlations (0.7 - 1, p < 0.05) from each VRP spike vaccine, focusing on IgG2a, functional assays, and Fc-gamma receptors FcgR3/R4 stimulated by the SARS-CoV-2 spike antigens (**Fig. 6A**). Strikingly, while numerous S1 and S2 correlates were identified, there was little overlap between the two. Recognition of S2 by various VRP spike sera was tied to Fc-effector mediated functions, while S1 demonstrated strong correlations with FcgR4, but was not statistically tied to effector functions (**Fig. 6A, B**). Using a peptide scanning array that spanned the majority of S2, we identified that heptad repeat region 2 (HR2), the fusion peptides (FP), and the stalk subregions drove much of the IgG2a recognition. Notably, we found that full spike, S2, and S2 subdomain-specific IgG2a and the phagocytic functional assays (ADNP/ADCP) had a strong correlation for the more distant heterologous VRP spike vaccines (HKU3, SARS-CoV, WIV1, SHC014). We also found that the VRP SARS-CoV-2 and the more protective spike vaccines (HKU3, RaTG13) IgG2a correlated with ADCD (**Fig. 6A**). Generally, FcR stimulation was clade-dependent and in some cases similar to the clade-dependent *in vivo* protection characterized above. FcgR4 was most activated by clade 1b and 2 VRP spike sera (SARS-CoV-2, RaTG13, and HKU3), and less so by clade 1a VRP spike sera (SARS, SHC014, and WIV1) (**Fig. 6B**). Consequently, we evaluated the capacity for the VRP spike serum to stimulate antibody-dependent cellular phagocytosis (ADCP) and neutrophil phagocytosis (ADNP) against the SARS-CoV-2 full spike as previously described at the univariate level (37, 55). Overall, we found notable and significant increases in ADNP (**Fig. 6C**) as well as significant ADCP against the SARS-CoV-2 full spike (**Fig. 6D**) in VRP spike vaccine sera. However, the magnitude of response was not clearly clade-dependent.

**Figure 6.**
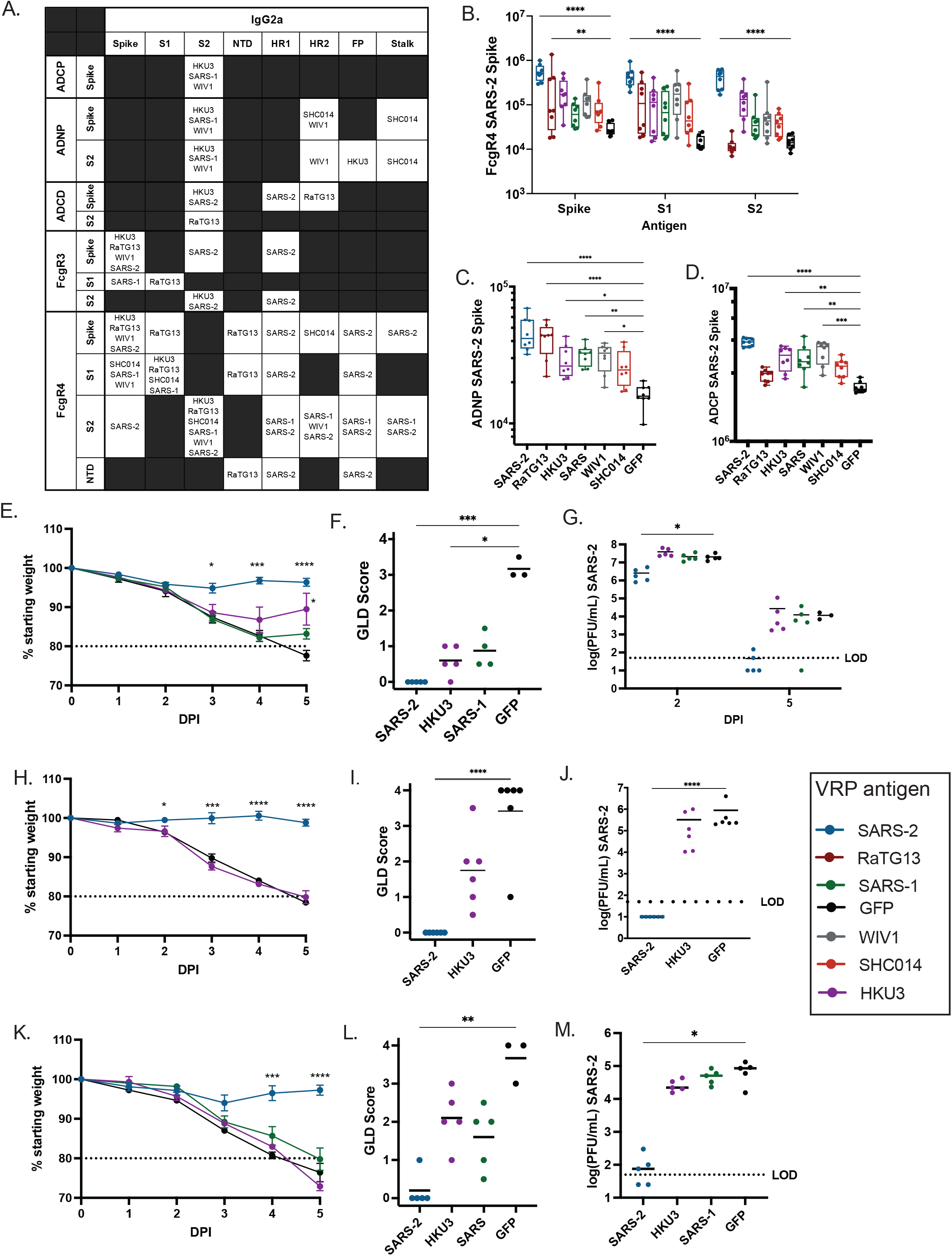
Non-neutralizing antibodies mediate protection *in vivo* through Fc function. **A.** Pearson’s correlation matrices were constructed of the systems serology assays for each VRP vaccination group (**Supplemental data**). VRP vaccine groups with strong correlations (0.7 – 1) between two assay results are listed in the table. **B.** FcgR4 stimulation against the SARS-2 spike (left), S1 (middle), and S2 (right) **C.** Phagocytic score of antibody-dependent neutrophil phagocytosis (ADNP) and **D.** Phagocytic score of antibody-dependent cellular phagocytosis (ADCP). **E.** body weights, **F.** GLD scores day 5 post infection, and **G.** SARS-2 lung titer in naïve mice after passive transfer of serum from vaccinated animals followed by intranasal infection of 10^4^ PFU SARS-2. **H.** Body weights, **I.** GLD scores day 5 post infection, and **J**. SARS-2 lung titer of young Fc receptor knockout BALB/c mice that were vaccinated with a sarbecovirus spike protein prior to infection with 10^4^ PFU SARS-2 intranasally. **K.** Body weights, **L.** GLD scores day 5 post infection, and **M.** SARS-CoV-2 lung titer of young Fc receptor knockout BALB/c mice that received prophylactic administration of serum from vaccinated wild-type BALB/c mice prior to infection with 10^3^ PFU SARS-CoV-2 intranasally. * p < 0.05, ** p <0.01, *** p < 0.001, **** p < 0.0001 after statistical testing described in methods. Body weights reported as percent of starting weight where horizontal line indicates 20% body weight lost and animal care humane endpoint. Titer samples that fell below the limit of detection (dotted line) were set to 25 PFU/mL.

Though we identified variable responses, our data as compiled by systems serology analysis demonstrate that VRP spike-specific, non-neutralizing antibodies stimulate FcR effector functions. These functions are mechanistically linked to FcgR4 binding IgG2a, and additional heatmap analysis of the heterologous VRP spike sera (**Fig. S7**) indicated that, not only did IgG2a cluster very well with itself, but the heptad repeat regions (HR1, HR2), and stalk subdomains of spike also exhibited the greatest strength of binding (**Fig. S7A**). When evaluating subclass binding within S2, a peptide scanning array also indicated that IgG2a bound with high significance to the HR1, HR2, fusion peptide (FP), and stalk subdomains of S2 (**Fig. S7B**).

Since HKU3 and RaTG13 spike vaccines showed the highest level of protection against heterologous SARS-CoV-2 challenge, we asked whether there was a correlation between domain specific antibody binding and disease severity. The magnitude of VRP HKU3 spike serum IgG2a binding the HR1 shared a very significant correlation (Pearson’s r = 0.92, p < 0.05, n = 4) with protection from disease as measured by AUC of body weight maintenance curves. Likewise, VRP RaTG13 sera S2 binding also showed a very significant correlation (Pearson’s r = 0.91, p < 0.05, n = 4) with protection from disease. Surprisingly, we found significant correlations between protection from disease and antibody binding in most S2 subdomains for WIV1 but in contrast to most other vaccine groups, failed to detect correlations between protection and functional assays like ADCD and ADCP for WIV1. Under further evaluation, we found that *in vivo* protection elicited by other VRP spike vaccines were linked to S2, but not single domains. Notably the SHC014 VRP spike vaccine was also closely linked to the NTD (**Fig. S7C**). These data suggest that the sum of smaller fractions of cross-reactive antibody responses may also contribute to protection in the context of the more distant, less protective vaccine strains in addition to the capacity of the binding antibodies to stimulate protective FcR effector responses.

These data implicate sequence conservation in linear epitopes of S2, especially HR1, in addition to cellular functional stimulation as a driver of the cross-protective, non-neutralizing antibody response elicited by VRP-vectored sarbecovirus spikes. Within S2, the HR2 subdomain is 100% conserved between the sarbecovirus spikes tested, while the HR1 subdomain contains sequence variation. The HR1 subdomain of RaTG13 and HKU3 is 100% and 98.7% identical to the SARS-CoV-2 HR1 subdomain, respectively. In contrast, the HR1 subdomains of the clade 1a sarbecovirus spikes which showed less protection, share 88.3-89.6% identity to the SARS-CoV-2 spike HR1 (**Fig. S7D**). These results support the hypothesis that the binding of non-neutralizing antibodies to conserved sequences within the HR1 domain may contribute to heterologous protection against SARS-CoV-2 challenge in our model.

### VRP spike vaccinations induce antibody-mediated protection via Fc effector mechanism

Given indications of non-neutralizing, antibody-dependent cellular function by serological assays, we conducted a prophylactic passive transfer experiment to further evaluate the role of antiserum in cross protection from clinical disease. Serum from VRP spike (SARS-CoV-2, SARS-CoV, HKU3, and GFP) vaccinated mice was pooled for a given group and then administered intraperitoneally into naïve mice (Taconic and Envigo, Envigo n = 5 reported). Twenty-four hours later, the mice were challenged with 10^4^ PFU of SARS-CoV-2 MA10 in a lethal challenge. Importantly, compared to GFP control, SARS-CoV-2 and HKU3 VRP serum recipients experienced significant reductions in weight loss (statistically significant by day 5 post infection), while the SARS-CoV VRP serum recipients developed more severe weight loss that was not significantly reduced compared to control serum recipients (**Fig. 6E**). We also observed significant reductions in GLD scoring by all groups, demonstrating that passive transfer of antibodies mitigated severe/lethal disease (**Fig. 6F**). This was in contrast with detected viral loads which were exclusively mitigated by VRP SARS-CoV-2 antigen (**Fig. 6G**). This provides further support to the hypothesis that neutralizing antibodies play a substantial role in limiting viral replication and disease, whereas non-neutralizing antibodies primarily mitigate disease pathology. Collectively, these data suggest that there is a clear role for antibodies, albeit non-neutralizing, in vaccine cross protection against SARS-CoV-2.

To further probe the mechanism of VRP spike cross-protection, we vaccinated FcR-deficient BALB/c mice (Taconic, n = 6) before lethal challenge with SARS-CoV-2 MA10. We found that, when Fc effector function is effectively eliminated, protection against SARS-CoV-2 disease in mice afforded by vaccination with VRP HKU3 spike was eliminated (**Fig. 6H–J**). Both rapid/sustained weight loss (**Fig 6H**) and GLD (**Fig. 6I**) were evident in the FcR KO mice immunized with VRP HKU3, and unlike WT mice, were statistically indistinguishable from GFP-vaccinated mice. However, FcR-deficient mice vaccinated with VRP SARS-CoV-2 spike were still protected from clinical disease and virus replication, again consistent with a strong homologous neutralizing antibody response (**Fig. 6J**). This indicates that cross protection is mechanistically linked to FcR-mediated responses despite a potent homologous protective profile.

To further evaluate the role for FcR effector function (e.g. macrophages, neutrophils) in VRP spike cross-protection *in vivo*, we conducted a prophylactic passive transfer experiment in FcR-deficient BALB/c mice (Taconic, n = 5), prior to lethal challenge with SARS-CoV-2 MA10. As evidenced by weight loss (**Fig. 6K**) and GLD scores (**Fig. 6L**), we found that the SARS-CoV-2 spike homologous sera still protected against severe disease in the absence of FcR effector function. Additionally, we found significant reductions in virus titer in the lungs of animals that received VRP SARS-CoV-2 spike sera, again supporting a strong homologous neutralizing antibody response (**Fig. 6M**). However, we found that the heterologous SARS-CoV and HKU3 VRP sera failed to protect in FcR-deficient mice (**Fig. 6L–N**). When compared to wild-type mice as previously described, FcR-deficient mice had increased GLD and increased weight loss, with no decrease in virus titer in the lungs. FcR-deficient mice that received heterologous VRP sera surpassed 20% weight loss on day 5 post infection, indicative of lethal disease. This indicates that strain-specific antibodies capable of neutralization can protect from disease, but cross protection is mechanistically linked to non-neutralizing, FcR-mediated responses.

Altogether, our data indicate that neutralizing antibodies can protect from disease, but are oftentimes limited to homologous challenges after VRP vaccination. Rather, non-neutralizing FcR function is a primary driver of VRP spike antibody-mediated cross protection from sarbecovirus disease.

## DISCUSSION

The emergence of SARS-CoV-2 underscores the tragic global consequences of a recently emerged zoonotic virus. The COVID-19 pandemic has resulted in massive human suffering and global economic upheavals with millions of deaths. When considering the continued spread of SARS-CoV-2 VOC, coupled with large numbers of zoonotic reservoir strains poised for cross species movement, robust countermeasures that elicit broad, cross-protective immune responses offer considerable hope for controlling sarbecovirus epidemics. Though a potent neutralizing antibody response is a benchmark for COVID-19 vaccine efficacy, recent work has identified that SARS-CoV-2 S2P mRNA vaccines elicited limited cross neutralizing antibody titers against heterologous sarbecoviruses and the SARS-CoV-2 Omicron VOC (18, 56, 57). Moreover, several studies have suggested a potential role for non-neutralizing antibody function in protection against SARS-CoV-2 disease (35, 36, 58), highlighting a critical need for identification of additional correlates associated with pan-sarbecovirus protection. As alphavirus replicons are under development as COVID-19 vaccines that induce mucosal, humoral and cellular immune responses, they provide innovative models for understanding cross immune mechanisms (15–17). In the present study, we found that non-neutralizing antibodies can contribute to broad cross-vaccine protective immunity across clade 1a, 1b, and clade 2 sarbecoviruses, with as little as 75% amino acid identity between spike protein amino acid sequences. Our studies further support an important protective role for non-neutralizing antibodies, especially in cases of heterologous virus infection across distant sarbecoviruses, via antibody interactions with FcR effector functions as a driver of protective immunity (36). In addition to T cell immunity (59), a good universal vaccine will likely stimulate multiple arms of the B cell-driven immune response – including potent type-specific, as well as broadly cross-neutralizing and non-neutralizing antibodies that promote FcR effector functions.

Consonant with a potent neutralizing antibody response described previously (44), VRP SARS-CoV-2 spike vaccination prevented severe disease in young mice and significantly reduced virus replication in the airway after SARS-CoV-2 MA10 challenge. Highly potent mRNA vaccines target neutralizing antibody responses to the S1 RBD domain (60, 61) and potent neutralizing antibodies targeting one or more epitopes in the RBD (28), NTD (62), or S2 (63, 64) have been identified in the spike protein. Consonant with established work, we found the homologous neutralizing antibody response stimulated by the VRP platform preferentially targeted the receptor-binding domain (targeted by VRP SARS-CoV-2 S) and/or the N-terminal domain (both targeted by VRP SHC014 S). S1 is highly divergent across sarbecoviruses, but contains conserved regions that stimulate cross-neutralizing antibodies (28, 34). However, we failed to detect protective levels of cross-neutralizing antibodies elicited by heterologous VRP spike vaccines after prime and boost, similar to other SARS-CoV-2 vaccines (18, 60). While this finding may be related to VRP vaccine dose, our data also suggest that different sarbecoviruses may focus neutralizing antibody responses to different sites within S1, potentially complicating universal vaccine platforms focused exclusively on the RBD, especially when applied to outbred populations, like the human population (65, 66).

Although heterologous spike vaccines failed to elicit cross neutralizing antibodies against SARS-CoV-2 and still allowed infection in the lungs, these vaccines reduced viral loads in the lungs and provided partial protection from SARS-CoV-2 disease in *in vivo*. A similar phenotype, mediated by coronavirus nucleocapsid-based vaccines that stimulate T cells, has been reported following SARS-CoV and MERS-CoV challenge (59). While our data does not explicitly rule out a contribution of T cells in mediating protection after VRP vaccination, passive antibody transfer, systems serology, and vaccine studies in wild-type versus FcR deficient mice mechanistically link these non-neutralizing antibody functions as strong drivers of cross protection

In our model, cross-protection was highly clade dependent, as clade 1b and clade 2 VRP spikes protected with higher efficacy against SARS-CoV-2 disease than clade 1a VRP spikes. Clade 2 HKU3 contains as much antigenic diversity with the SARS-CoV-2 spike as clade 1a WIV1. However, the VRP HKU3 spike elicited near-complete protection from SARS-CoV-2 MA10 disease while the VRP WIV1 spike did not. This clade-dependent protection suggests that specific domain conservation, rather than overall sequence homology, drives the development of protective antibodies in our model. Additionally, we found that VRP SARS-CoV-2 spike vaccination, while unable to prevent infection, provided partial protection against disease induction by the heterologous bat virus, HKU3-SRBD MA. This result aligns with current data regarding spike-based vaccine efficacy against SARS-CoV-2 variants in the human population; where currently approved vaccines may not prevent variant infection in all cases but significantly reduce disease severity and death (67–70). Our model noted no cross protection following contemporary human coronavirus spike vaccinations against SARS-CoV-2. While contrary to some earlier correlative studies (22, 71–74), these differences may reflect repeat group 1A/2B human ß-coronavirus infections, which might result in more cross protective humoral responses, highlighting an area of future investigation. Overall, our results indicate that when faced with future sarbecovirus emergence events, cross protection from vaccine-mediated SARS-CoV-2 immunity has the potential to reduce disease severity or breadth of transmission. Still, this protection is less likely to extend beyond the sarbecovirus subgenus.

There is increasing evidence of a potential role for non-neutralizing antibodies in long-term SARS-CoV-2 vaccine protection, especially against VOC (39, 58, 75). Consistent with these data, while heterologous antibodies elicited by VRP 3526-delivered sarbecovirus spikes did not neutralize SARS-CoV-2, we detected and characterized cross-protective non-neutralizing antibody activity through passive transfer experiments and systems serology. Our data highlights non-neutralizing antibodies as correlates of heterologous vaccine-mediated protection in an FcR-dependent manner. Furthermore, we detected strong FcR stimulation by antibodies that recognized the SARS-CoV-2 spike as well as the S1 and S2 regions, especially the heptad repeat 1 (HR1) region of S2, from the heterologous vaccines. Sarbecovirus strain variation-dependent diversity of response in addition to the many instances of correlated stimulus supports the hypothesis that the non-neutralizing cross protection may also be a function of the cumulative effect of epitope conservation rather than driven by single or few broadly cross-protective epitopes that are conserved across β-coronaviruses (76). Thus, Fc-effector immunity may depend upon the number and quality of cross binding epitopes as well as the abundance of cross-reactive antibodies, coupled with the types of FcR/effector mediated phenotypes.

Further supporting our results, we found that both homologous and heterologous protection was less robust in aged animals, consistent with existing work (44, 51). Notably, age related waning of FcR effector functions is thought to impact both vaccine efficacy and infection response (77–79). Notably, neutrophil responses wane in aged populations and become dysregulated after pulmonary infection (79). Additionally, overall effector cell function (e.g. infected cell killing) becomes impaired with increased age (77, 78). Thus, as a function of increasing virus challenge dose in aged animals, increased VRP sarbecovirus spike vaccine failure is consistent with reduced FcR mediated protection, a prospect that will need to be carefully investigated in future vaccine studies focused on the elderly.

While this work has shown that the VRP platform is a valuable experimental platform for studying cross-protective coronavirus immunity, especially regarding non-neutralizing antibody responses, the study design also highlights many areas for further investigation. The same susceptibility loci appear to regulate sarbecovirus pathogenesis in mice and humans (53), and the mouse model reproduces key aspects of acute and chronic SARS-CoV-2 induced disease (42, 43). Furthermore, mouse models of SARS-CoV-2 disease have proven to be robust platforms for predicting SARS-CoV-2 vaccine performance in humans (53, 80–82) and the alphavirus replicon strategy has shown utility as a vaccine platform (83–86). However, systems serology responses following alphavirus vaccination in humans have not been reported, including any reporting on non-neutralizing antibody functional activity. This study is the first that clearly implicated FcR-mediated protection following alphavirus VRP vaccination in any species as well as the first to directly correlate the FcR mechanism of cross-protection to disease outcome. However, our data was drawn from a limited sample size for correlation to disease (n = 4), prompting a need for further mechanistic investigation. Still, detailed systems serology studies have also suggested a robust correlative role for FcR-mediated protection after mRNA vaccination and the durability of protection when compared to prior infection (39). Additionally, FcR mechanisms of protection have been implicated in HIV vaccination (45), influenza vaccination (87–89), and DENV prior infection (90).

Although speculative, the results identified here may likely be relevant to understanding the mechanisms that promote vaccine-induced SARS-CoV-2 immunity in humans, as also evidenced by the fact that a recombinant SARS-CoV-2 RBD protein vaccine conferred cross protection in the absence of a potent cross neutralizing antibody response (91). However, most vaccine designs have not been tailored to maximize protective FcR effector functions despite clear animal model studies that have demonstrated that specific activation of distinct FcγR-mediated pathways significantly improves antibody-mediated protection as well as sustained and robust immune responses (39, 92–94). In addition, the exact antibodies and epitopes that contribute to cross protection via FcR mediated activities remain unclear, but obviously would help guide future pan-coronavirus therapeutic development including both monoclonal antibody treatment and vaccines. This is especially interesting in the unique case of cross-protection we observed with VRP HKU3 S. Investigating epitope conservation, in this case, may identify novel spike epitopes that contribute to broad cross-protection. For example, our work would propose that a pan-sarbecovirus vaccine would benefit from inclusion of the HR1 spike subdomain to induce cross-protective FcR effector responses. These studies also suggest that the VRP platform represents a valuable system both for human vaccine delivery and for dissecting the aspects of vaccine-induced immunity that mediate protection. It will be essential to determine whether these results extend beyond standard inbred mouse strains by testing their impact in outbred populations, such as the Collaborative Cross (95, 96), or other models of SARS-CoV-2-induced disease like non-human primates to further probe the mechanisms of cross protection discussed.

As SARS-CoV-2 is the second sarbecovirus to emerge in the 21^st^ century, other coronaviruses will likely arise in the future, including those with similar or different spike sequences to those examined in this study (e.g. SHC014 and WIV1) (4) and others (e.g. swine acute diarrhea syndrome (SADS) coronavirus) (97). Therefore, the inclusion of non-neutralizing, cross-protective epitopes informed by the results of our study may shift vaccine development toward a more comprehensive, cross-protective formulation that prevents life-threatening sarbecovirus disease and provide new insights for vaccine design against other highly heterogeneous RNA virus families, including *Coronaviridae*.

## FIGURES

**Figure S1. Pairwise sequence alignments of sarbecovirus spike proteins to the SARS-CoV-2 spike protein.** Global alignments constructed with Blosum62 matrix with free end gaps in Geneious Prime. Alignments of SARS-CoV-2 to **A.** RaTG13 **B.** HKU3 **C.** SARS **D.** WIV1 and **E.** SHC014. Blue – NTD, red – RBD, green – S2.

**Figure S2. VRP-vectored endemic coronavirus spike protein vaccinations do not protect against severe SARS-CoV-2 disease in young BALB/c mice. A.** Body weights and **B**. lung titer calculated for animals vaccinated with spike proteins from contemporary human CoVs. **C.** Percent survival of animals vaccinated with each spike protein. * p < 0.05, ** p <0.01, *** p < 0.001, **** p < 0.0001.

**Figure S3. Lung cytokine signatures in VRP-vaccinated mice.** Measured by BioPlex, cytokine signatures associated with immune responses were measured on days 2 and 5 post-infection with SARS-CoV-2 MA10. * p < 0.05, ** p <0.01, *** p < 0.001, **** p < 0.0001.

**Figure S4. VRP-vectored cross-protection wanes in aged animals with high dose lethal challenge.** Old mice (12 months) were challenged with 10^4^ PFU SARS-CoV-2 MA10 intranasally. **A.** Body weights calculated through the duration of the experiment on animals vaccinated with sarbecovirus spike proteins. Reported as percent of starting weight. **B.** Semi-quantitative macroscopic lung discoloration scoring upon tissue harvest. **C.** SARS-CoV-2 lung titer calculated via plaque assay. * p < 0.05, ** p <0.01, *** p < 0.001, **** p < 0.0001. Body weights reported as percent of starting weight where horizontal line indicates 20% body weight lost and animal care humane endpoint. Titer samples that fell below the limit of detection (dotted line) were set to 25 PFU/mL.

**Figure S5. Systems serology reveals novel mechanisms of protection.** The geometric mean fluorescent intensity (gMFI) value of each systems serology assay for a given serum sample plotted on a heatmap after log transformation.

**Figure S6. Correlation matrix of all systems serology metrics tested.** Pearson’s correlation coefficient calculated for each pair of systems serology assay for the listed set of post-boost serum samples. **A.** Full data set of metrics for all samples. **B.** VRP GFP **C.** VRP SARS-CoV-2 spike **D.** VRP RaTG13 spike **E.** VRP HKU3 spike **F.** VRP SARS-CoV spike **G.** VRP WIV1 spike **H.** VRP SHC014 spike post boost serum samples tested.

**Figure S7. Identification of likely cross-protection drivers. A.** Heatmap identifying IgG2a peptide recognition clusters. **B.** Background-subtracted IgG2a peptide binding within S2 **C.** VRP vaccine groups with strong correlations between disease (AUC bodyweight curve) and antibody factors as identified via Pearson’s correlation matrices. **D.** Multiple sequence alignments of S2 subdomains.

## METHODS

### Biosafety and institutional approval

All experiments were conducted after approval from the UNC Chapel Hill Institutional Biosafety Committee and Institutional Animal Care and Use Committee according to guidelines outlined by the Association for the Assessment and Accreditation of Laboratory Animal Care and the US Department of Agriculture. All vaccinations were performed at ABSL2 while all infections and downstream assays were performed at ABSL3 in accordance with Environmental Health and Safety. All work was performed with approved standard operating procedures and safety conditions for SARS-CoV-2. Our institutional ABSL3 facilities have been designed to conform to the safety requirements recommended by Biosafety in Microbiological and Biomedical Laboratories (BMBL), the US Department of Health and Human Services, the Public Health Service, the Centers for Disease Control and Prevention (CDC), and the National Institutes of Health (NIH). Laboratory safety plans have been submitted, and the facility has been approved for use by the UNC Department of Environmental Health and Safety (EHS) and the CDC.

### Cell Lines and viruses

All cell lines and viruses were confirmed mycoplasma-negative. All viruses used were subjected to next-generation sequencing prior to use. Vero E6 cells were maintained in Dulbecco’s Modified Eagle’s Medium (DMEM) supplemented with 5% FBS and anti/anti. Baby Hamster Kidney (BHK21) cells were maintained in α-MEM supplemented with 10% fetal bovine serum, L-glutamine, and 10% tryptose phosphate broth. Mouse adapted SARS-CoV-2 MA10(42) and SARS-CoV-2 nanoLuciferase reporter viruses were developed based on the SARS-CoV-2 WA1 reference strain (98) and propagated from a cDNA molecular clone as previously described. Mouse adapted bat virus HKU3 was generated from a cDNA molecular clone (11, 53) and mutations were inserted that cause pathogenesis in mice. To generate the SARS-CoV-2 spike domain panel, the backbone sequence from bat virus SHC014 (4) was used. SHC014 spike sequences were replaced with corresponding fragments of the sequence encoding SARS-CoV-2 spike segments (RBD, RBD+, NTD, RBD-NTD, S1) and viruses were generated from the cDNA clone.

### VEE VRP3526 vaccine preparation

The sequence encoding CoV spike proteins (below) were cloned into the pVR21 vector containing Venezuelan equine encephalitis virus strain 3526 non-structural proteins. RNA from template pVR21 constructs and VEE 3526 helper constructs encoding glycoprotein and capsid proteins was transcribed using Invitrogen T7 mMessage mMachine *in vitro* transcription kit. Purified RNA was electroporated into BHK21 cells in the ratio of 2:1:1 pVR21 construct: VEE 3526 glycoprotein: capsid. Supernatant was harvested and purified 24 hours post-electroporation and ultracentrifuge concentrated through sucrose cushion. VRP titers were determined through immunofluorescent staining to detect VEE-associated proteins. All VRPs were confirmed to not cause cytopathic effect in cell culture before administration to mice. Spike sequences used to construct VRP vaccines:

|  |  |  |  |
| --- | --- | --- | --- |
| SARS-CoV-2 | MT020880.1 | HKU3 | FJ211859.1 |
| RaTG13 | MN996532.2 | NL63 | AY567487.2 |
| SARS-CoV | AY278741 | OC43 | UDM84911.1 |
| SHC014 | KC881005.1 | 229E | KY621348.1 |
| WIV1 | KC881007.1 | HKU1 | HM034837.1 |

### Mice, vaccination, and infection

BALB/cAnNHsd were obtained from Envigo (strain 047) and delivered at either 8-10 weeks (young) or 11-12 months (old) for vaccination and housed in groups under standard conditions. Mice were vaccinated with 2×10^4^ VRP in a 10 μl phosphate-buffered saline footpad inoculation and boosted with the same dose 3 weeks post-prime. Baseline, pre-boost, and pre-challenge serum was collected via submandibular bleed. Four weeks post-boost, mice were infected with 10^3^ or 10^4^ (where specified) PFU SARS-CoV-2 MA10 or 10^5^ PFU HKU3 MA-SRBD in 50 μl PBS intranasally under ketamine-xylasine anesthesia. The challenge doses were at least one log higher than the suspect infectious dose in humans (99). For adoptive and passive transfer experiments, 200 μl serum from vaccinated mice was transferred to naïve mice via intraperitoneal injection 24 hours prior to challenge. Mice were weighed daily through the course of infection, and a subset’s respiratory function was tracked daily using whole body plethysmography (48). Mice were euthanized at 2 and 5 days post infection via isoflurane overdose. FcR-knockout mice were obtained from Taconic (Model 584), on a BALB/cAnNTac background delivered at 6-10 weeks old due to strain availability. The relevant infectious challenge dose for this strain was determined to be 10^3^ PFU SARS-CoV-2 MA10 due to strain-specificities, and mice were infected and monitored as described above.

### Mouse tissue collection and analysis

After euthanasia, blood was collected into phase separation tubes by cardiocentesis or severing the vena cava and allowed to clot before centrifugation to separate serum. Lungs were scorerd for gross discoloration, indicating congestion and/or hemorrhage, based on a semi-quantitative scale of mild to severe discoloration covering 0 to 100% of the lung surface. The left lung was collected and injected with 10% neutral buffered formalin to expand airways before storage in fixative for 7 days before histopathological processing. Of the right lung lobes, the inferior lobe was collected in ~1 mL TRIzol reagent with glass beads and the superior lobe was collected in ~1 mL phosphate buffered saline with glass beads. Both inferior and superior lobes were homogenized in a MagnaLyser and debris was pelleted. Virus in the lungs was quantified from the superior lobe via plaque assay. Briefly, virus was serial diluted and inoculated onto confluent monolayers of Vero E6 cells, followed by agarose overlay. Plaques were visualized on day 2 post infection via staining with neutral red dye. Lung cytokines were quantified from the superior lobe using the Bio-Plex Pro Mouse Cytokine 23-Plex Immunoassay. RNA from the inferior lobe was reserved for additional downstream assays.

### Neutralization assays

A serial dilution (1:20 initially, followed by a 3-fold dilution) of pre-challenge serum was incubated in a 1:1 ratio with SARS-CoV-2-nLuc (98) to result in 800 PFU virus per well. Serum virus complexes were incubated at 37C with 5% CO2 for 1 hour. Following incubation, serum virus complexes were added to a confluent monolayer of Vero E6 cells and incubated for 48 hours at 37C with 5% CO_2_. After incubation, luciferase activity was measured with the Nano-Glo Luciferase Assay System (Promega) according to the manufacturer specifications. Neutralization titers (EC50) were defined as the dilution at which a 50% reduction in RLU was observed relative to the virus (no antibody) control.

### Systems Serology

SARS-CoV-2 and other sarbecovirus and control antigens were resuspended in water to a final concentration of 0.5 mg/mL and linked to magnetic Luminex beads (Luminex Corp, TX, USA) through carbodiimide NHS ester linkages. Specific antigens were coupled to individual bead regions. Biotinylation of antigens were done using the NHS-Sulfo-LC-LC kit, and excess biotin was removed using Zebra-Spin desalting and size exclusion columns. Antigen coupled beads were then incubated with serum at various dilutions (1:100 for IgG2a, IgG2b, IgG3, IgM, 1:200 for IgG1, and 1:750 for Fcγ-receptor binding) in a 384-well plate (Greiner, Germany) overnight at 4°C. Unbound material was washed and detection of isotypes and subclasses were done using PE-conjugated anti-IgG1, -IgG2a, -IgG2b, -IgG3, -IgM. PE-Streptavidin (Agilent Technologies, CA, USA) was coupled to recombinant and biotinylated mouse FcγR2b, FcγR3, and FcγR4A at a 1:1000 dilution. Secondary detection was done at room temperature for 1 hour, and unbound material was removed by washing. Relative binding per antigen was determined on an IQue Screener PLUS cytometer (IntelliCyt).

Antibody-dependent cellular phagocytosis (ADCP) and neutrophil phagocytosis (ADNP) assays were done as previously described (100). Mouse serum was incubated with cultured monocytes or primary neutrophils at a concentration of 1:100 on preformed immune complexes on fluorescent neutravidin microspheres. Cells were fixed with 4% paraformaldehyde (PFA) and identified by gating on microsphere-positive cells. Phagocytic score was quantified by the (percentage of microsphere-positive cells) x (MFI of microsphere-positive cells) divided by 100000. Antibody-dependent complement deposition (ADCD) was done as previously described (101). Relative complement deposition was quantified through flow cytometry, as measured by fluorescein-conjugated goat IgG that targets the guinea pig complement C3b.

Correlations between Fab, FcR, and functional assays were done using GraphPad Prism using Spearman’s coefficients. Statistical significance, as defined by p < 0.05, was corrected for multiple comparisons using Benjamini-Hochberg correction. Other analyses such as PLS-DA were done on R using the systemsseRology pipeline available on GitHub (GitHub - LoosC/systemsseRology: Machine learning tools (for the analysis of systems serology data) are also available. Each assay contained pre-immune and post-vaccination sera, as well as PBS controls to account for batch effects. All other calculations are described below.

Key reagents used in systems serology assays

|  |  |  |
| --- | --- | --- |
| Anti-mouse IgG1-PE | Southern Biotech | 1144-09 |
| Anti-mouse IgG2a-PE | Southern Biotech | 1155-09L |
| Anti-mouse IgG2b-PE | Southern Biotech | 1186-09L |
| Anti-mouse IgG3-PE | Southern Biotech | 1191-09L |
| Anti-mouse IgM-PE | Southern Biotech | 1021-09 |
| Anti-CD66b Pac Blue | BioLegend | 305112 |
| Anti-CD3 | BD Biosciences | 558117 |
| Anti-CD16 | BD Biosciences | 557758 |
| Anti-CCL4 | BD Biosciences | 550078 |
| Anti-C3b | MP Biomed | 855385 |
| SARS-CoV-2 WT Spike | Sino Biological | 40589-V08H4 |
| SARS-CoV-2 D614G Spike | Sino Biological | 40589-V08H8 |
| SARS-CoV-2 WT S1 Domain | Sino Biological | 40591-V08H |
| SARS-CoV-2 WT Receptor Binding Domain (RBD) | Sino Biological | 40592-V08H |
| SARS-CoV-2 WT S2 Domain | Sino Biological | 40590-V08B |
| SARS-CoV-2 WT N-terminal Domain | Sino Biological | 40591-V49H |
| SARS-CoV-2 Alpha Variant S | Sino Biological | 40589-V08B6 |
| SARS-CoV-2 Beta Variant S | Sino Biological | 40589-V08B7 |
| SARS-CoV-2 Delta Variant S | Sino Biological | 40589-V08B16 |
| SARS-CoV-2 Omicron Variant S | Sino Biological | 40589-V08H26 |
| Human Coronavirus OC43 S | Sino Biological | 40607-V08B |
| Human CoV HKU1 S (isolate N5) | Sino Biological | 40606-V08B |
| Human Cytomegalovirus (HCMV) Glycoprotein B (gB) | Sino Biological | 10202-V08H1 |
| Human Coronavirus 229E Spike | Sino Biological | 40605-V08B |
| SARS-CoV-1 Spike | Sino Biological | 40634-V08B |
| PE-Streptavidin | Agilent Technologies | PB32-10 |
| Guinea Pig Complement | Cedarlane | CL4051 |
| Protein Transport Inhibitor | BD Biosciences | 554724 |
| Brefeldin-A | Sigma | B7651 |
| NHS-Sulfo-LC-LC Kit | ThermoFisher | 21435 |
| Zebra-Spin Desalting and Chromatography Columns | ThermoFisher | 89882 |
| Fix & Perm Cell Permeabilization Kit | ThermoFisher | GAS002S-100 |
| R Studio V 1.4.1103 | RStudio, PBC | Open Source |
| GraphPad Prism | GraphPad Software, LLC | Ragon Site License |
| FlowJo V. 10.8 | FlowJo, LLC | www.flowjo.com/so |
|  |  | lutions/flowjo/down loads |
| iQue Forecyt | Sartorius | 60028 |
| iQue Screener Plus | Intellicyt/Sartorius | 11811 |
| 384-well HydroSpeed Plate Washer | Tecan | 30190112 |
| MagPlex Microspheres | Luminex MFG | MC12001-01<br>(Cataloged by region) |
| Green Fluorescent Neutravidin Microspheres | ThermoFisher | F8776 |
| Red Fluorescent Neutravidin Microspheres | ThermoFisher | F8775 |

### Enzyme-Linked Immunosorbent Assay

#### Full-length spike protein ELISA titer

All serum samples tested by ELISA assay were heat-inactivated at 56°C for 30 min to reduce risk from possible residual virus in serum. ELISA binding titer for full-length spike protein was measured as described before (102). Essentially, full-length spike protein at 2 μg/mL in Tris Buffered Saline (TBS) pH 7.4 was coated in the 96-well microtiter plate for 1 hour at 37°C. The wells were blocked with 3% milk in TBS containing 0.05% Tween 20 (TBST) for 1 hour, then serially diluted serum samples were added (1:100 - 1:24,300) to the wells and incubated for an additional hour at 37°C. The plate was washed three times using wash buffer (TBS containing 0.2% Tween 20), then respective goat anti-mouse IgG (Catalog # A16072), IgG1 (Catalog # PA1-74421), or IgG2A (Catalog # M32207) was added at 1:2000 and incubated for 1 hour at 37°C. The plate was washed three times using wash buffer, then 3,3’,5,5’-Tetramethylbenzidine (TMB) Liquid Substrate (Sigma-Aldrich) was added to the plate, and absorbance was measured at 450 nm using a plate reader (Molecular Devices SpectraMax ABS Plus Absorbance ELISA Microplate Reader) after stopping the reaction with 1 N HCl.

#### RBD or NTD ELISA titer

All serum samples tested by ELISA assay were heat-inactivated at 56°C for 30 min to reduce risk from possible residual virus in serum. ELISA binding titer for Spike RBD or NTD was measured as described above with minor modifications. 96-well microtiter plate was coated with Streptavidin (Invitrogen) at 4 μg/mL in TBS pH 7.4 for 1 hour at 37°C. The wells were blocked with 1:1 Non-Animal Protein-BLOCKER^™^ (G-Biosciences) in TBS for 1 hour. Biotinylated spike RBD or NTD antigen (1 μg/ml) was captured onto the streptavidin-coated wells, then serially diluted serum samples (1:100 - 1:24,300) were added to the wells and incubated for 1 hour at 37°C. The plate was washed three times using wash buffer (TBS containing 0.2% Tween 20), then goat anti-mouse IgG (Catalog # A16072) was added at 1:2000 and incubated for 1 hour at 37°C. The plate was washed three times using wash buffer, then 3,3’,5,5’-Tetramethylbenzidine (TMB) Liquid Substrate (Sigma-Aldrich) was added to the plate, and absorbance was measured at 450 nm using a plate reader (Molecular Devices SpectraMax ABS Plus Absorbance ELISA Microplate Reader) after stopping the reaction with 1 N HCl.

### Statistical testing

All statistical analyses were conducted in GraphPad PRISM 9. To assess the statistical significance of weight loss, significance was calculated by two-way ANOVA comparing each spike vaccinated group to the GFP control. In cases where mortality was observed, significance was calculated via Mixed-effects analysis The significance of virus and antibody titers was calculated via one-way ANOVA comparing each spike vaccinated group to the GFP control. To assess the significance of lung discoloration and histopathological lung damage scoring, significance was calculated via Brown-Forsythe and Welch’s ANOVA. In all cases, testing was corrected for multiple comparisons using Dunnett’s multiple comparisons test, Dunnett’s T3 test when total samples <50. Significance reported as * p < 0.05, ** p <0.01, *** p < 0.001, **** p < 0.0001.

## ACKNOWLEDGEMENTS

GA and the Systems Serology Lab at Ragon are supported by Mark and Lisa Schwartz, Terry and Susan Ragon, and the SAMANA Kay MGH Research Scholars award. GA and the Systems Serology Lab also receives funding from the Massachusetts Consortium on Pathogen Readiness (MassCPR), the Gates Global Health Vaccine Accelerator Platform, and the NIH (3R37AI080289-11S1, R01AI146785, U19AI42790-01, U19AI135995-02, U19AI42790-01, P01AI165072, U01CA260476 – 01, CIVIC75N93019C00052). VKB receives funding from NIH grant K01OD026529. RSB and MTH receive funding from NIH grant P01AI158571. RSB receives funding from NIH/NCI North Carolina Seronet Center for Excellence grant U54 CA260543. LEA is supported in part by NIH 2T32AI007419-26 Molecular Biology of Viral Diseases Predoctoral Training Grant. This project was also supported in part by the North Carolina Policy Collaboratory at the University of North Carolina at Chapel Hill with funding from the North Carolina Coronavirus Relief Fund established and appropriated by the North Carolina General Assembly. We would like to also thank Sharon Taft-Benz for valuable input coordinating, directing, and managing studies in ABSL3 facilities.

