## Supplementary material for "Fc mediated pan-sarbecovirus protection after alphavirus vector vaccination": Adams et al Supplement.pdf

### S1. Pairwise sequence alignments of Sarbecovirus spike proteins to the SARS-2 spike protein

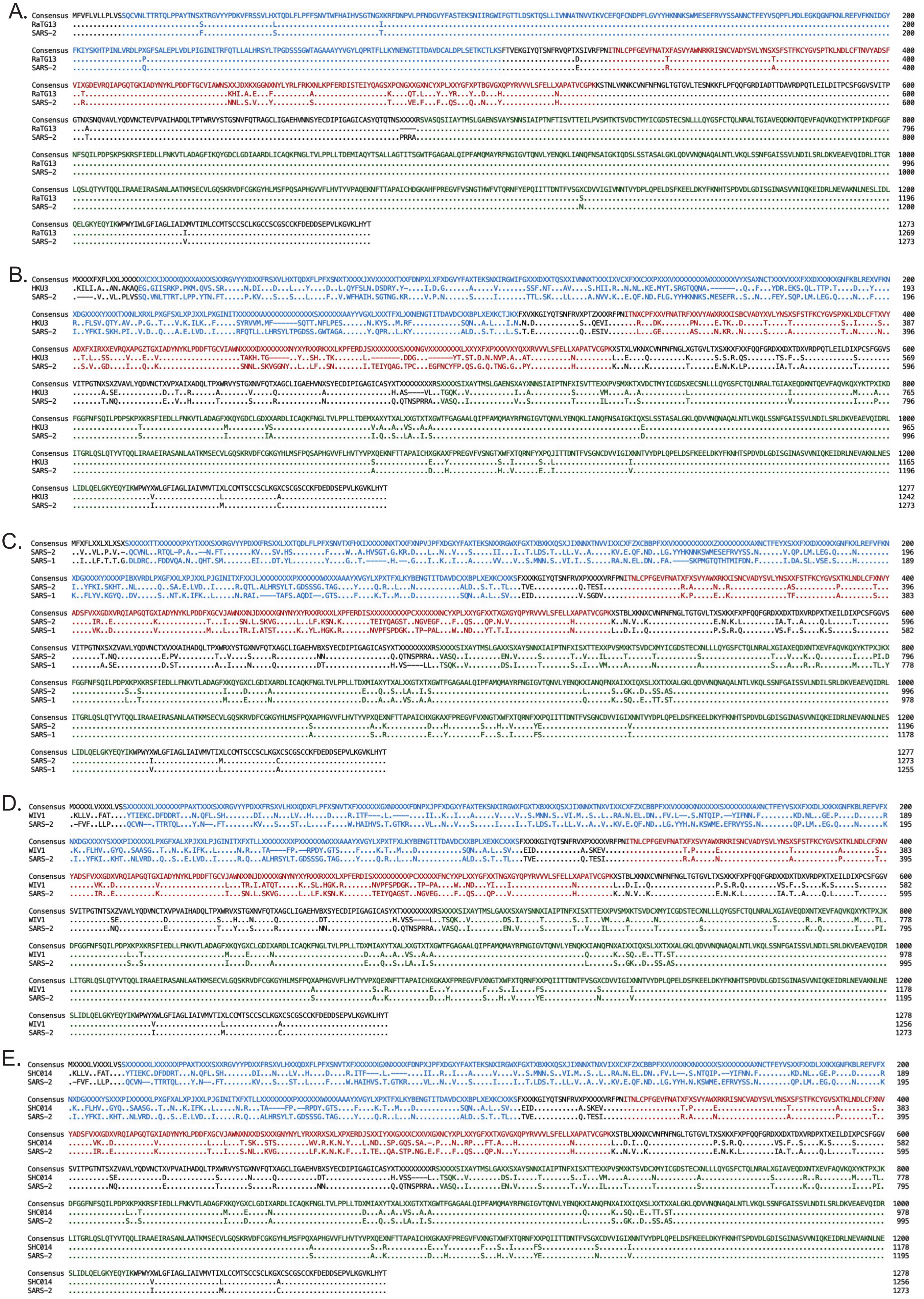

#### S2. VRP-vectored endemic CoV spike vaccinations do not protect against SARS-2 disease

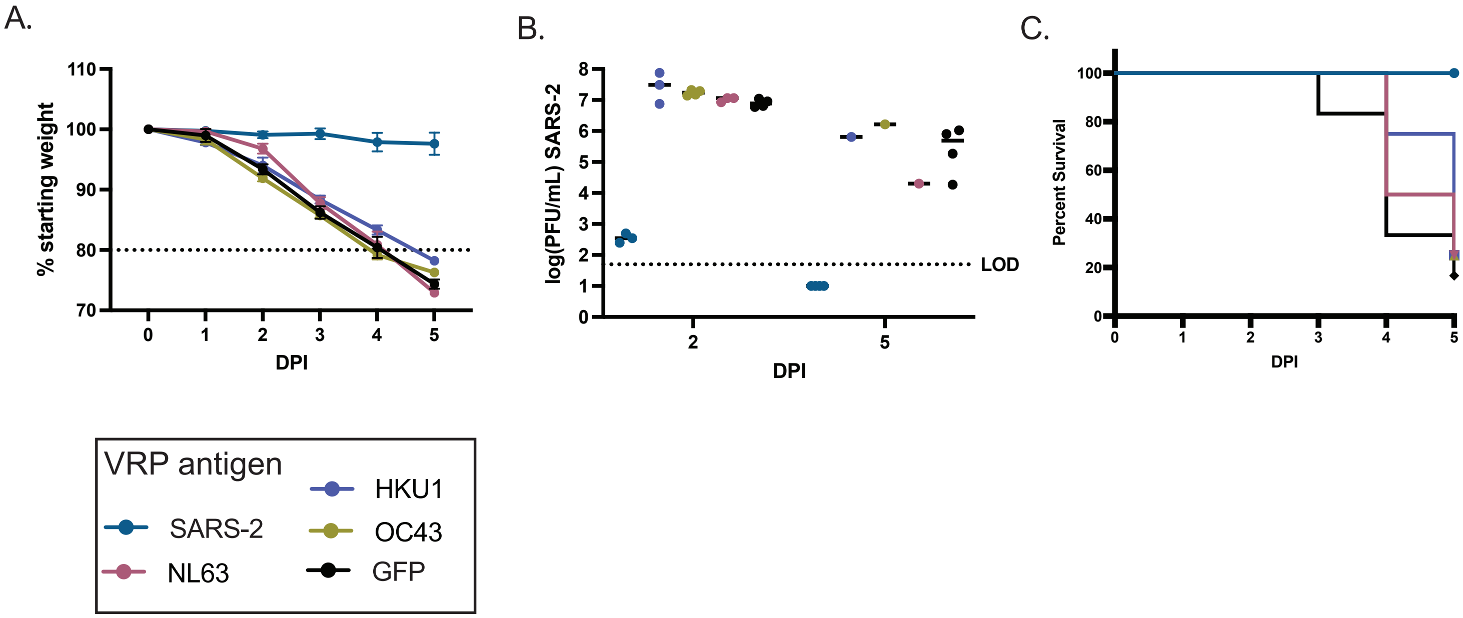

#### S3. Lung cytokine signatures in VRP-vaccinated mice.

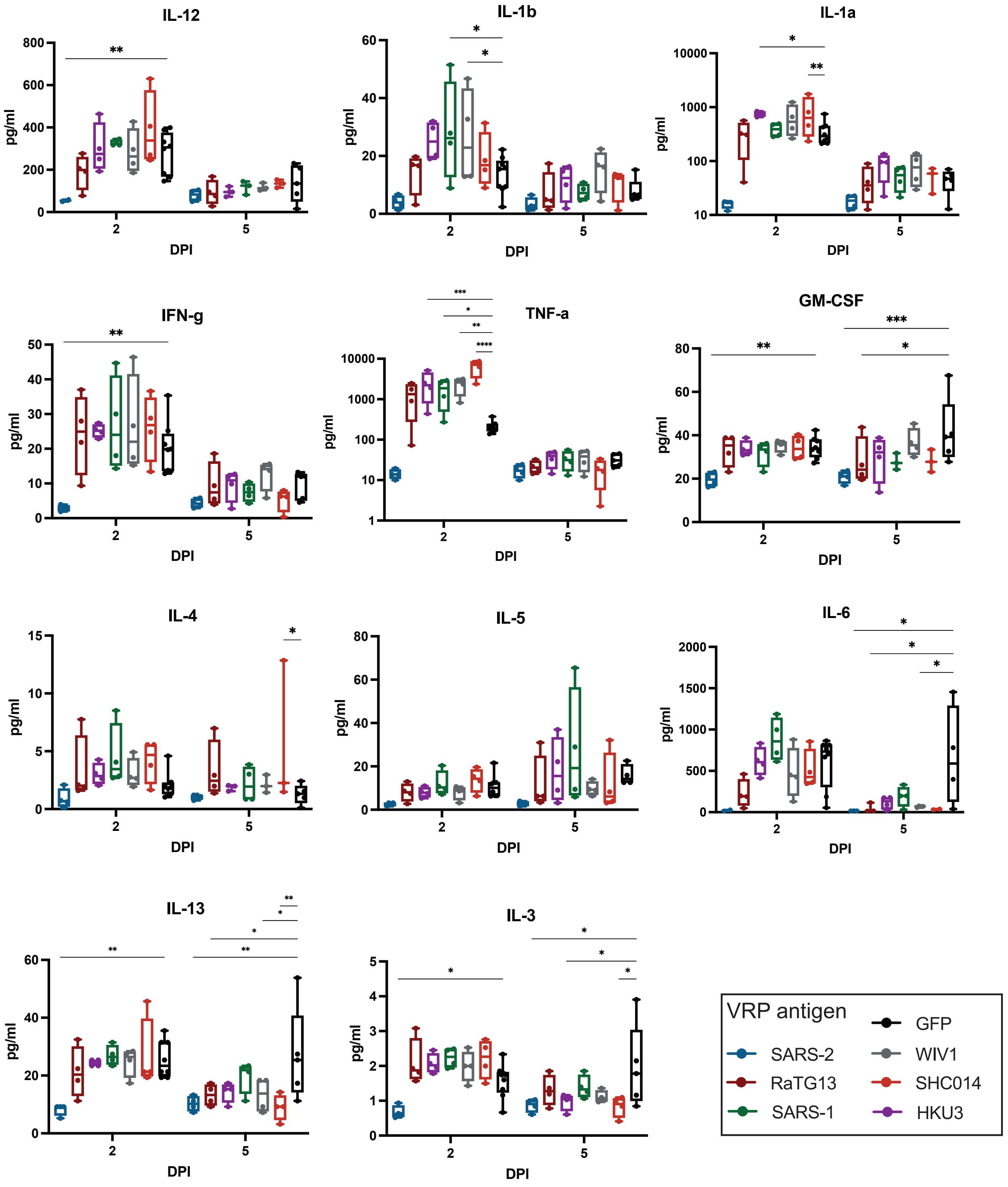

#### S4. VRP-vectored cross-protection wanes in aged animals with high-dose lethal challenge

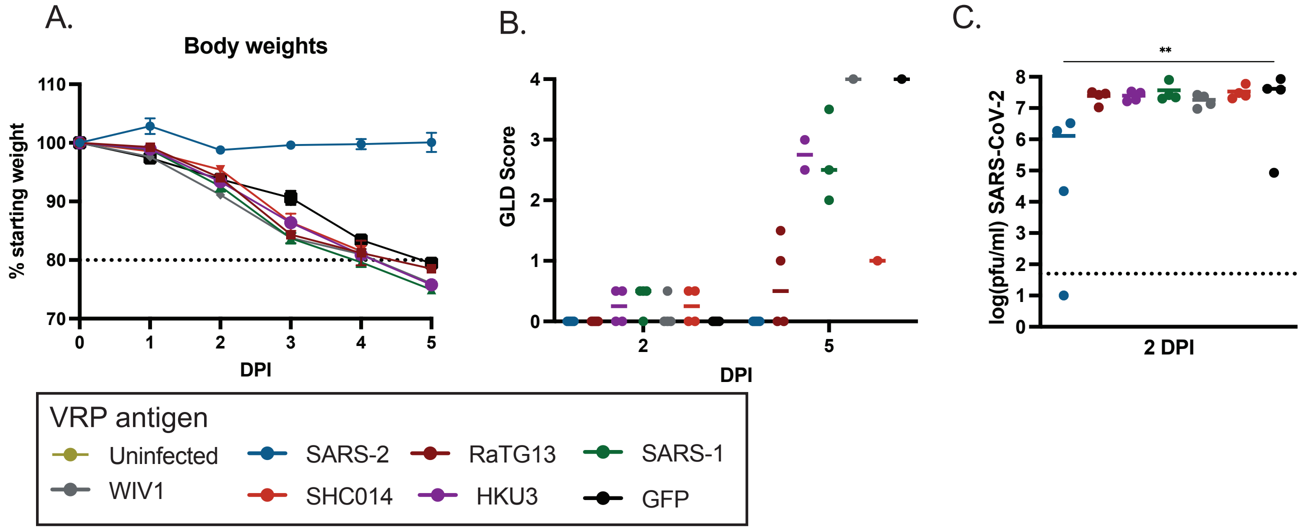

#### S5. Systems serology reveals novel mechanisms of protection

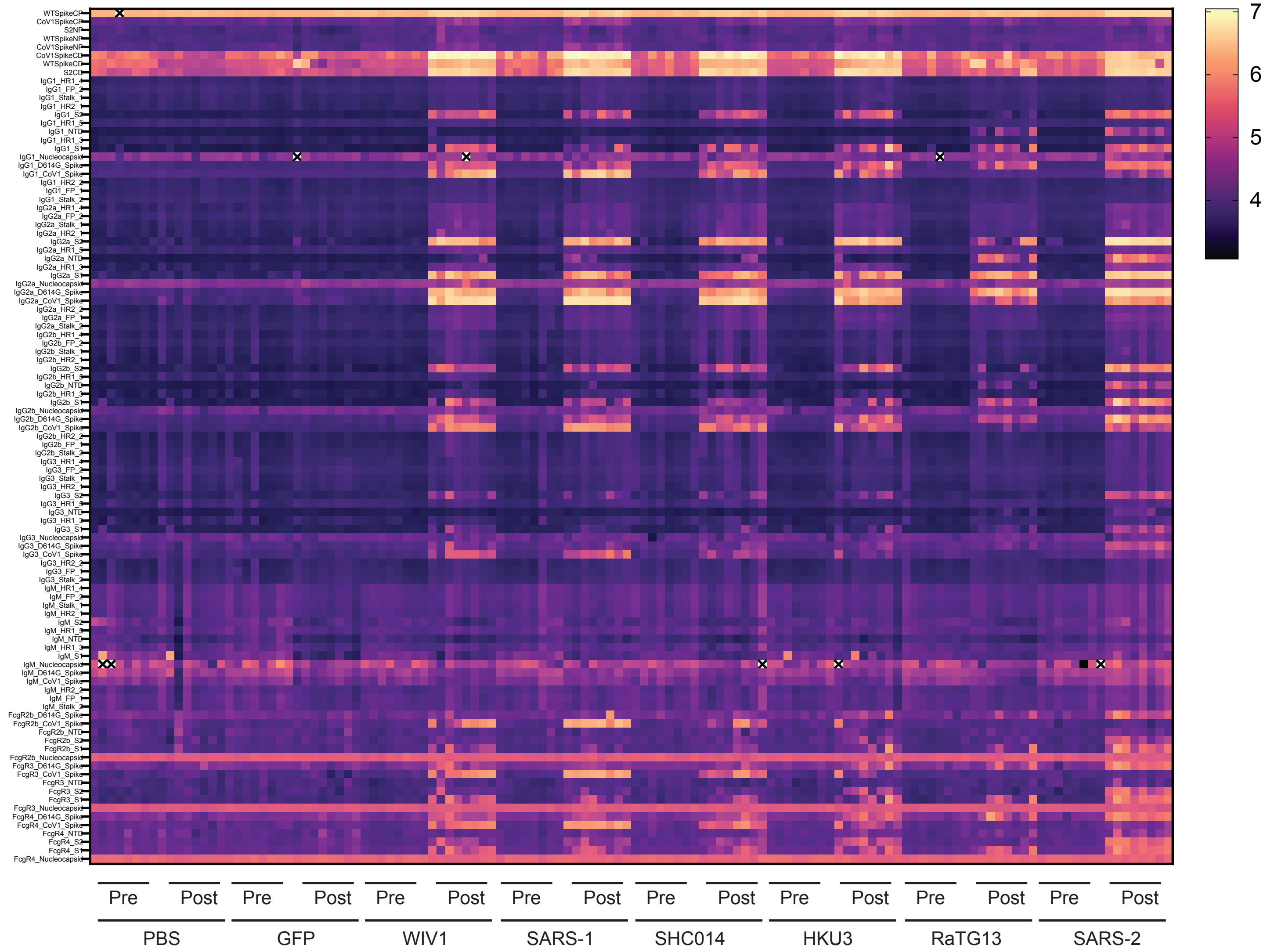

### S6. Correlation matrices of all systems serology metrics for all samples tested

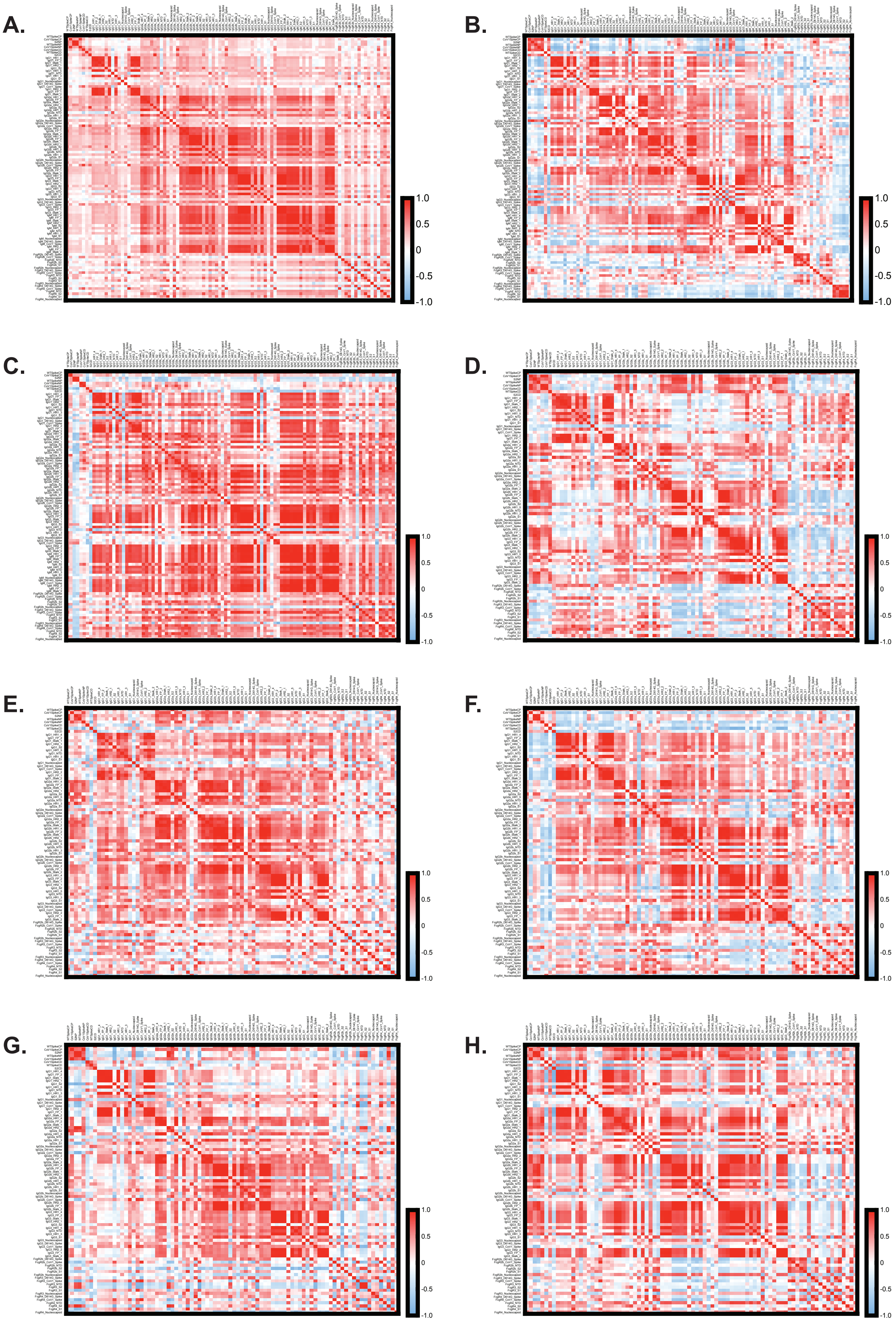

#### S7. Identification of likely cross-protection drivers

**A.**

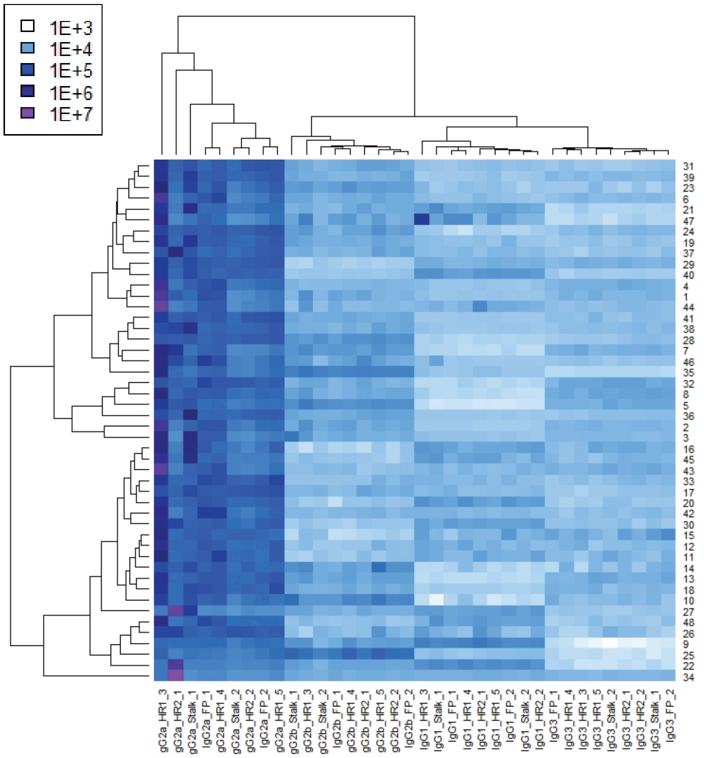


**B.**

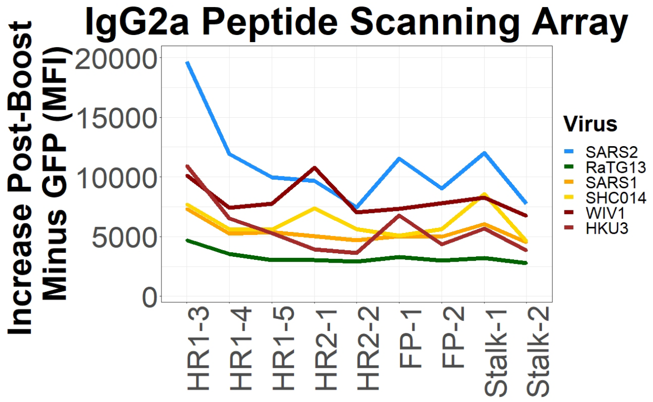


**C.**

| | IgG2a | | | | | | | | ADCD | ADNP | | ADCP |
|---|---|---|---|---|---|---|---|---|---|---|---|---|
| | Spike | S1 | S2 | NTD | HR1 | HR2 | FP | Stalk | S2 | Spike | S2 | Spike |
| **AUC % Starting Weight** | | | HKU3 RaTG13 | HKU3 SHC014 | HKU3 WIV1 | HKU3 WIV1 | HKU3 WIV1 | HKU3 WIV1 | HKU3 RaTG13 SHC014 SARS-2 | HKU3 | HKU3 | HKU3 RaTG13 SARS-1 |

**D.**

```
            FP1   FP2

Consensus   DPXKPTKRSFIEDLLFNKVTLADAGFM

SARS-2      ..S..S....................I   834
HKU3        ..S........................   803
RaTG13      ..S..S....................I   830
SARS-1      ..L........................   816
SHC014      ..L........................   817
WIV1        ..L........................   817
```

```
            HR2-1   HR2-2

Consensus   VDLGDISGINASVVNIQKEIDRLNEVAKNLNESLIDLQELGKYEQYIK

SARS-2      ................................................   1211
HKU3        ................................................   1180
RaTG13      ................................................   1207
SARS-1      ................................................   1193
SHC014      ................................................   1194
WIV1        ................................................   1194
```

```
            HR1-3   HR1-4   HR1-5

Consensus   GVTQNVLYENQKXIANQFNXAIXXIQESLXXTXXALGKLQDVVNQNAQALNTLVKQLSSNFGAISSVLNDILSRLDKVEAEVQ

SARS-2      ............L......S..GK..D..SS.AS.................................................   992
HKU3        ............L......S..GK.....SS.AS.................................................   961
RaTG13      ............L......S..GK..D..SS.AS.................................................   988
SARS-1      ............Q......K..SQ.....TT.ST.................................................   974
SHC014      ............Q......K..SQ.....TT.ST.................................................   975
WIV1        ............Q......K..SQ.....TT.ST.................................................   975
```

```
            Stalk-1   Stalk-2

Consensus   NTVYDPLQPELDSFKEELDKYFKNHTSPDVDLGDISGINASVVNIQKEIDRLNEVAKNLNESLIDLQELGKYEQYIKWPWYVWLGFIAGLIAIVMVTILL

SARS-2      .................................................................................I................M.   1234
HKU3        ....................................................................................................   1203
RaTG13      .................................................................................I...........I....M.   1230
SARS-1      ....................................................................................................   1216
SHC014      ....................................................................................................   1217
WIV1        ....................................................................................................   1217
```
